# Evolutionary responses to acquiring a multidrug resistance plasmid are dominated by metabolic functions across diverse *Escherichia coli* lineages

**DOI:** 10.1101/2022.07.22.501110

**Authors:** Laura Carrilero, Steven Dunn, Alan McNally, Michael Brockhurst

**Author notes:** Equal contribution (Laura Carrilero, Steven Dunn).

## Abstract

Multidrug resistance (MDR) plasmids drive the spread of antibiotic resistance between bacterial lineages. The immediate impact of MDR plasmid acquisition on fitness and cellular processes varies among bacterial lineages, but how the evolutionary processes enabling the genomic integration of MDR plasmids vary is less well understood, particularly in clinical pathogens. Using diverse *Escherichia coli* lineages experimentally evolved for ∼700 generations, we show that the evolutionary response to gaining the MDR plasmid pLL35 was dominated by chromosomal mutations affecting metabolic and regulatory functions, with both strain-specific and shared mutational targets. The expression of several of these functions, such as anaerobic metabolism, is known to be altered upon acquisition of pLL35. Interactions with resident mobile genetic elements, notably several IS-elements, potentiated parallel mutations, including insertions upstream of *hns* that were associated with its upregulation and the downregulation of the plasmid-encoded extended-spectrum beta-lactamase gene. Plasmid parallel mutations targeted conjugation-related genes, whose expression was also commonly downregulated in evolved clones. Beyond their role in horizontal gene transfer, plasmids can be an important selective force shaping the evolution of bacterial chromosomes and core cellular functions.

## Introduction

*Escherichia coli* is a common cause of human and animal infections. In particular, multidrug resistant (MDR) lineages cause serious invasive diseases and pose significant challenges for effective treatment (Mathers et al., 2015). The emergence of these MDR lineages is typically associated with the acquisition of one or more multidrug resistance plasmids (Dunn et al., 2019). Such plasmids often encode resistances against multiple antibiotic classes, including clinically important frontline treatments (Cantón and Coque, 2006). In particular, the acquisition and dissemination of extended spectrum beta-lactamase (ESBL) genes in *E. coli* is largely mediated by the transfer of MDR plasmids (Dunn et al., 2019). The ESBL CTX-M-15 confers resistance to cephalosporins and has been widely disseminated by ISEcp1, which serves to both mobilise this gene among plasmids, and increase its expression by replacing its native promoter (Lartigue et al., 2006). The abundance and diversity of ESBL plasmids in *E. coli* makes controlling their spread incredibly challenging. Understanding the factors that determine the successful integration of these plasmids into bacterial genomes is therefore a high priority.

The response of bacterial cells to plasmid acquisition can vary extensively between bacterial lineages. For example, multiple studies report that the immediate growth or fitness impact of acquiring an identical plasmid ranges from negative to positive between different lineages of a species (Alonso-del Valle et al., 2021; Dunn et al., 2021; Gama et al., 2020). These differences have been linked to gene content variation among the bacterial genomes in some cases (Alonso-del Valle et al., 2021). In addition, the transcriptional response to plasmid acquisition can also vary between bacterial lineages in terms of the identity of differentially expressed chromosomal genes, the numbers of bacterial genes affected, and the magnitude of their change in expression level (Dunn et al., 2021). Collectively, these comparative experimental data suggest that there will be lineage-specific or perhaps even genotype-specific responses to MDR plasmid acquisition that are contingent upon genetic variation among the recipient bacterial genomes.

In lineages where plasmid acquisition imposes high fitness costs and/or causes appreciable disruption to cellular function(s), we would expect plasmid carriage to be rare because plasmid-bearing cells will be selected against in antibiotic-free environments, where the benefits of plasmid encoded traits do not outweigh the fitness cost of plasmid carriage (Bergstrom et al., 2000; Harrison and Brockhurst, 2012; San Millan, 2018; San Millan and Maclean, 2017).

However, an emerging theme in evolutionary studies of plasmid-host interactions is that plasmid acquisition often acts as a catalyst for evolutionary changes in the bacterial chromosome and/or the plasmid itself, which enable the stable genomic integration of newly acquired plasmid replicon(s) even when these are at first costly (Brockhurst and Harrison, 2022). Such evolutionary responses to plasmid acquisition can take the form of compensatory mutation(s) that resolve specific genetic conflicts between plasmid and chromosomal genes to ameliorate the cellular disruption these cause (Hall et al., 2021; Loftie-Eaton et al., 2017; Millan et al., 2015; Porse et al., 2016). Alternatively, these evolutionary responses can be more extensive, involving coadaptation of the chromosome and plasmid, whereby multiple mutations occurring on both replicons are necessary to assimilate the plasmid into the genome (Bottery et al., 2019, 2017; Jordt et al., 2020). To date, few studies have compared the evolutionary responses of diverse lineages to acquiring a new plasmid at the genomic level (Benz and Hall, 2022; Jordt et al., 2020; Porse et al., 2016). As such little is known about how the evolutionary processes of genomic integration of an MDR plasmid varies between genomically diverse bacterial lineages.

Here we take a comparative experimental evolution approach to test how the evolutionary response to acquisition of the MDR plasmid pLL35 varies between genomically diverse lineages of *E. coli*. pLL35 was obtained from a *Klebsiella* clinical isolate and encodes multiple antibiotic resistance genes, including the extended-spectrum beta-lactamase CTX-M-15 (Dunn et al., 2021). Replicate populations of five *E. coli* strains, including both environmental and clinical isolates and the lab strain MG1655, carrying pLL35 were serially-passaged with or without cefotaxime for ∼700 bacterial generations, alongside plasmid-free controls propagated without cefotaxime. Bacterial population density and plasmid frequency was monitored over time by selective plating and colony PCR. At the end of the experiment, we compared the growth and cefotaxime resistance of evolved clones and obtained their whole genome sequences and, for MG1655, the ancestral and evolved transcriptomes. The growth response to selection varied between lineages and according to cefotaxime treatment, but no change in cefotaxime resistance was observed between treatments. We observed parallel mutations in evolved plasmid-carriers at loci or within operons associated with a range of functions including cellular metabolism, regulation of MGEs, and plasmid conjugation.

## Results

### Growth kinetic and resistance responses to selection

To test for the initial effect of plasmid acquisition on bacterial growth we compared growth kinetic parameters for each of the ancestral *E. coli* strains with or without pLL35. Acquisition of pLL35 reduced growth in the ancestral *E. coli* strains (Figure S1; statistical tables provided in Table S1) indicating that plasmid carriage imposed a fitness cost in all strain genetic backgrounds used here. Despite this initial fitness cost of the plasmid, however, we observed no appreciable plasmid loss in any of the plasmid-containing populations during the ∼700 generations selection experiment either with or without cefotaxime (Figure S2), indicating that pLL35 was stably maintained regardless of positive selection for the encoded CTX-M-15 extended-spectrum beta-lactamase.

We next quantified the change in growth kinetic parameters and cefotaxime resistance for a randomly chosen evolved clone per replicate population from the end of the selection experiment relative to their ancestor (these evolved clones were also used in subsequent genome sequencing). The response of growth kinetic parameters to selection varied among strains and between treatments (Figure 1; statistical tables provided in Table S2). Notably, the strains MG1655 and F022 showed higher growth rates and maximum densities relative to their ancestors than did strains EL39, F104 and F054. Moreover, plasmid bearing clones evolved with cefotaxime tended to show higher performance across a range of growth kinetic parameters than evolved plasmid-free control clones. However, the level of resistance to cefotaxime in evolved plasmid carriers did not vary with strain or cefotaxime selection (ART ANOVA strain F_4,60_ = 1.4161 P = 0.24; cefotaxime selection F_2,60_ = 2.8666 P = 0.06), although high variability was observed among evolved clones for some strains, suggestive of divergence between replicate populations (Figure S3).

**Figure 1.**
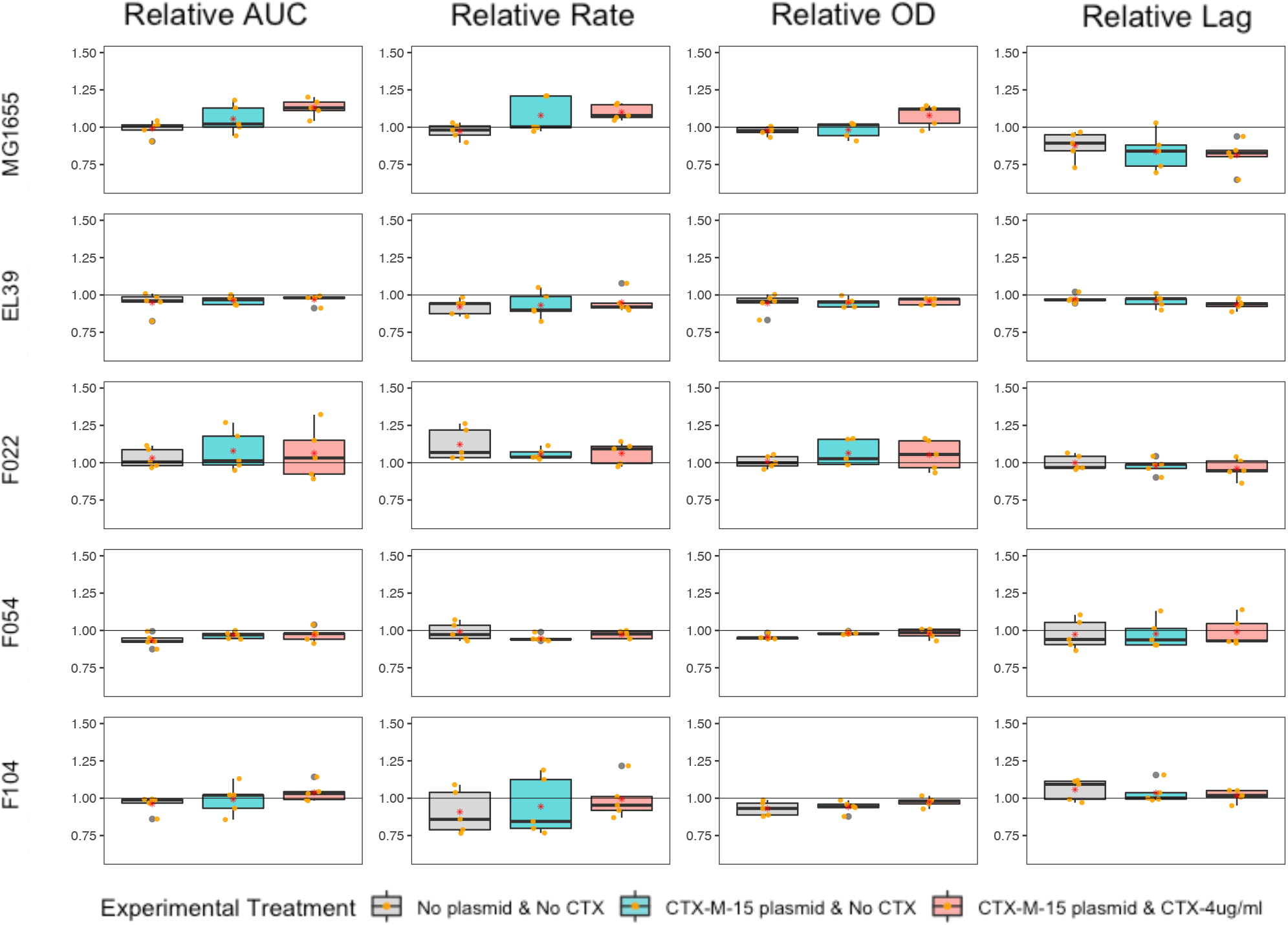
Growth kinetics of evolved bacterial clones relative to their ancestor. Boxplots represent the change in growth kinetic parameters for evolved clones relative to their ancestor. Each strain is shown in a separate row and each growth kinetic parameter is shown in a different column, as indicated by the labels. Each evolution treatment is denoted by a colour (grey, plasmid-free control, C; red, plasmid-carrier without cefotaxime, EP; blue plasmid-carrier with cefotaxime, EX). Datapoints show the mean of technical replicates for each individual evolved clone.

### Genetic responses to selection

To determine the genetic response to selection and how this varied among strains and treatments we obtained whole genome sequences for a randomly chosen evolved clone per replicate population. Evolved clones had gained between 0-45 mutations, including mutations located both on the chromosome (range = 4 – 43 mutations) and the plasmid (range = 0 - 21 mutations) in the evolved plasmid-carrying clones. 4-out-of-50 evolved plasmid-carrying clones were hypermutators, having acquired mutations in mismatch repair, whilst none of the evolved clones from control populations were hypermutators. Excluding hypermutator clones, of all the observed SNVs, 12.1% (n=23) were synonymous, 22.2% (n=42) were intergenic, and the remaining (n=124) 65.6% were nonsynonymous. There were also 18 instances of IS movement and 4 deletions. The number of nonsynonymous mutations per evolved clone did not vary among *E. coli* strains (P=0.0931 ANOVA) or between treatments (P=0.6603 ANOVA). Next, to distinguish mutations putatively associated with an evolved response to plasmid-acquisition, we identified the subset of chromosomal loci acquiring mutations that were exclusive to plasmid-carriers (i.e., loci that never acquired mutations in the corresponding plasmid-free controls) for further analysis (Figure 2). This subset contained 14 loci that were mutated in more than one independently evolved clone. Such parallel evolution is suggestive of selection having acted upon the mutations at these loci, which were then analysed further. Functional analysis of these loci revealed enrichment for transcriptional regulators (rpoS, putA, norR, nhaR, fadR) and inorganic ion transport and metabolism (putP, mgtA, artP, nanT), with the remaining loci representing singleton COG categories.

**Figure 2.**
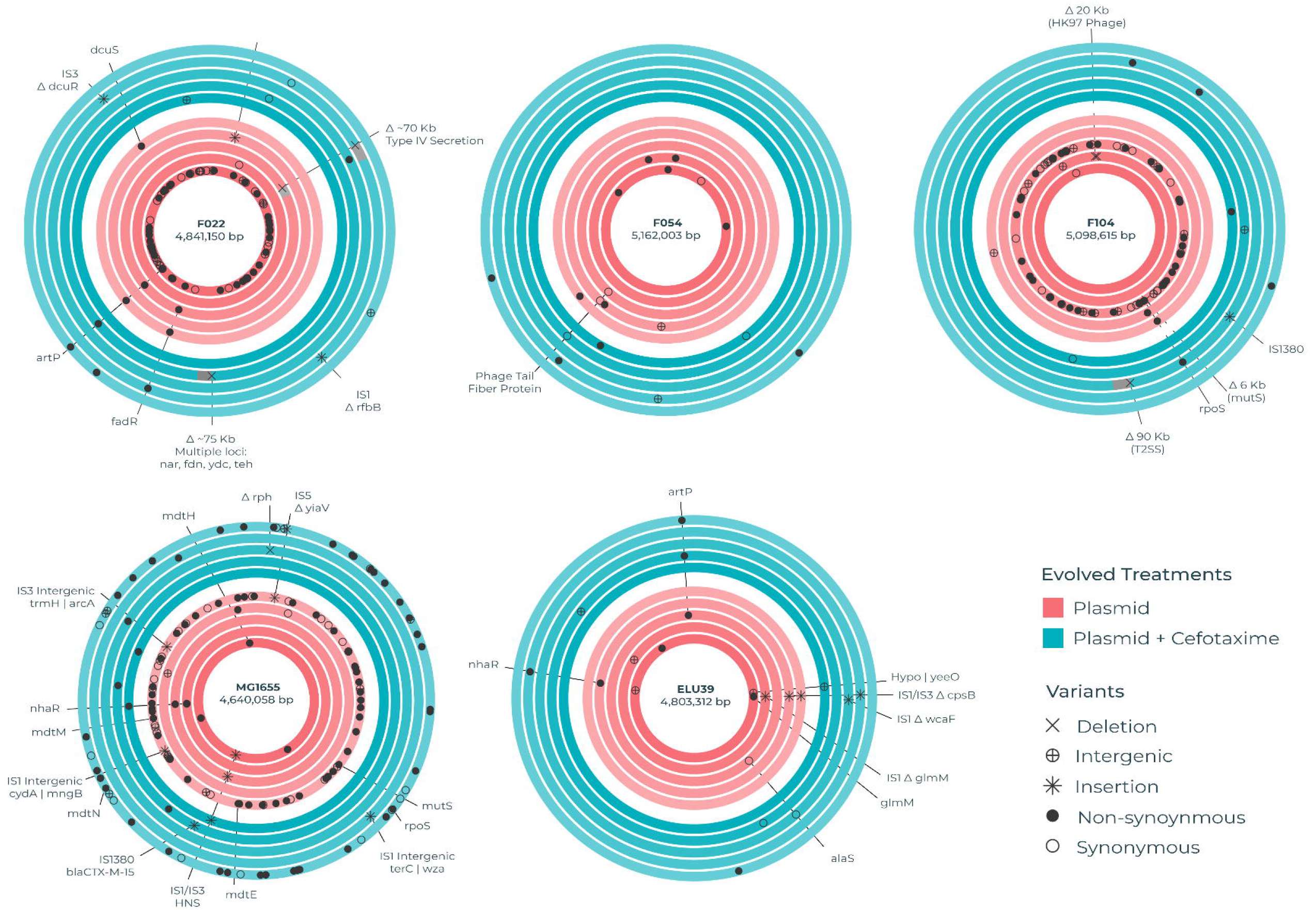
Chromosomal loci targeted by mutations in evolved plasmid-carriers. Each circular track represents the chromosome of an independently evolved clone. A single randomly chosen clone was genome sequenced per evolving line. As such the five replicate evolved clones per strain (as labelled) per treatment (Blue for plasmid-carriers evolved with cefotaxime, EX; Red for plasmid-carriers evolved without cefotaxime, EP) are shown as concentric tracks. Loci that acquired mutations during evolution in plasmid-carriers but not in the corresponding plasmid-free control are shown by markers denoting the type of mutation (see visual key). Loci that acquired parallel mutations in multiple independently evolving lines per strain per treatment have been labelled with the corresponding gene name or locus tag.

At a number of chromosomal loci and operons we observed parallel evolution occurring between strains (∼22% of SNVs, n=78; Figure 2, S4), suggesting a common evolved response to the plasmid among the divergent *E. coli* genetic backgrounds. The most commonly mutated locus was the arginine transporter gene artP, with mutations affecting 9 evolved clones across multiple replicates of F022 and ELU39 both with and without cefotaxime selection, and a single replicate of F054. Several of these mutations were clustered within a similar region of the ArtP protein, which encodes the main ABC transporter domain and may therefore affect arginine transport. Other transporters affected by mutations in multiple strains were the sialic acid transporter gene nanT and magnesium transporting ATPase gene mgtA. Several other metabolic functions also displayed between strain parallelism. These included key components of anaerobic metabolism including fatty acid metabolism and anaerobic respiration: fadR a multifunctional regulator of fatty acid metabolism, was mutated in 4 evolved clones of strains ELU39 and F104 where mutations clustered between amino acid residues 29-35. In both cases, these residues occur in the turn between an alpha helix and beta strand of a putative HTH *gntR* type conserved domain, which may therefore have some transcriptional impact. In F104 a variant was detected in *fadI* from the same operon which catalyses the final step in fatty acid oxidation. Mutations were observed in *norR* (A418A, A448V), an anaerobic nitric oxide reductase transcriptional regulator, in both MG1655 and F022. The sodium/proline metabolism *put* operon also exhibited between strain parallelism, with *putP* containing an identical D55G mutation across two replicates of F022, and a single MG1655 isolate. In the F022 isolates, *putA* gained mutations at two separate amino acid residues (L213P, Y1073H). Other functions displaying between strain parallelism included: the transcriptional regulator *nhaR* controlling expression of the NhaA Na+/H+ antiporter protein in strains MG1655 and ELU39; the stress response sigma factor *rpoS* in strains MG1655 and F104; the mismatch repair gene *mutS* resulting in hypermutability in 4 evolved plasmid-carrying clones of strains MG1655, F104, and F022; and the glycerol metabolism operon *glp*.

In contrast, at several other loci we observed parallel evolution occurring only within evolved lines from a single strain, suggesting certain strain specific evolutionary responses to plasmid acquisition. In MG1655 several genes in the multidrug resistance operon *mdt* contained non-synonymous mutations, though no change in resistance to cefotaxime was detected in the clones carrying these mutations. Mutations were also observed in multiple MG1655 replicates in the *ydh* operon, including the monooxygenase *ydhR* which is involved in the metabolism of aromatic compounds, as well as ABC transporter *yjj*. Other examples of within strain parallelism affected hypothetical genes without known functions. Of the ∼78% of SNVs occurring as singletons amongst replicates, the majority related to transcriptional control or metabolic functions.

We observed multiple insertion sequence mediated mutations in evolved clones, suggesting that this was an important mutational mode for some strains in our study, although the propensity for IS-mediated mutations varied substantially among the *E. coli* strains. Whereas multiple IS-mediated mutations were observed in evolved clones of MG1655 (n=8), F022 (n=3), and ELU39 (n=5), only a single IS-mediated mutation occurred in F104, and none were observed in F054. In MG1655, we observed a common insertion of two separate insertion sequences (IS1 & IS3) upstream of *hns*, and thymidine kinase gene *tdk*, all within a 33 bp window [IS1 EPC = 2,587,180, IS1 EXA = 2,587,192, IS3 EPA = 2,587,213] in 3 evolved clones (with and without cefotaxime selection). The 26 amino acids adjacent to *hns* were unaffected, suggesting that the hns promoter is likely intact, but the transposon may form a discretely transcribed unit. In F104 and MG1655 we observed movement of IS1380, originating on pLL35 and encompassing the CTX-M-15 gene. In MG1655 the entire transposon integrated successfully into the chromosome, but in F104 the CTX-M gene was truncated. The successful duplication of IS1380 in MG1655 does not appear to have altered the cefotaxime MIC of the evolved clone. In ELU39 we observed a common truncation of Mannose-1-phosphate guanylyltransferase gene *cpsB*, mediated by two different IS elements, and an IS-element insertion affecting the Phosphoglucosamine mutase protein GlmM that does not interrupt the protein, but rather occupies the sequence immediately adjacent. Across two different replicates of ELU39, we observed an insertion of IS3 in the two-component regulatory element *dcuR*, and a non-synonymous mutation in the second component of that system, *dcuS*. This two component regulator controls anaerobic fumarate metabolism, and also weakly regulates the fumarate transporter encoded by *dctA*, in which we observed an insertion of IS1 causing truncation of this gene. This suggests multiple routes likely to alter fumarate uptake and metabolism, and alongside mutations targeting other aspects of anaerobic metabolism across multiple strains (e.g., fatty acid metabolism and anaerobic respiration), indicates that anaerobic metabolism was a key target of selection in plasmid-carriers.

Few mutations were observed in the pLL35 plasmid sequence in the evolved clones, with observed variants frequently targeting genes involved in plasmid conjugation (Figure S5). Two evolved clones of F054 contained pLL35 with nonsynonymous mutations affecting *traD* encoding the coupling protein and *traI* encoding the multifunctional conjugation protein, and an evolved clone of MG1655 contained pLL35 with a nonsynonymous mutation affecting *traI*. In F104 an evolved clone from the cefotaxime selected treatment contained pLL35 with a complete deletion of the conjugational machinery that rendered the plasmid conjugation deficient. Excluding this deletion, all other replicates were able to conjugate. Additionally, we observed that pLL35 from an evolved MG1655 had acquired an IS-element inserted intergenically between a hypothetical protein and putative fimbrial subunit gene *flmA*.

Parallel mutations accounted for ∼22% (n=78) of SNVs across our dataset, with the remaining 78% occurring as singletons amongst replicates. Of these genes, the majority also related to transcriptional control or metabolic functions.

### Transcriptional responses to selection in MG1655

Given that MG1655 showed the strongest phenotypic responses to selection and contained a combination of strain-specific and shared mutational targets in evolved plasmid carriers, hereafter we focused our analyses on better understanding the evolutionary responses of this strain across treatments and replicate populations. We performed RNAseq on the ancestral genotype with or without pLL35 and on each of the evolved clones. In the ancestral MG1655, acquisition of pLL35 caused a total of 17 chromosomal genes to be moderately differentially expressed (Log_2_ fold change ≥ 1, FDR ≤0.05), with functions primarily related to metabolism (e.g. *dcuB, fumB, malK, gadX, hycD*; Figure S6). Three genes in the L-threonine degredation operon *tdc* showed a common signature of upregulation. This modest transcriptional impact of the plasmid is consistent with the small but significant reduction in ancestral growth we observed in this strain, and more broadly is similar to the scale of transcriptional disruption caused by pLL35 in the other strains previously reported, albeit affecting largely different genes.

Next, we analysed differential expression in the evolved clones relative to their ancestor to detect evolved transcriptional responses. A large fraction of significantly differentially transcribed genes (Log_2_ fold change ≥ 1.5, FDR ≤0.05) were common to all treatments, that is occurring in both evolved plasmid-free and evolved plasmid-containing clones (n = 1450 genes; 59%), presumably representing general adaptation to the lab environment. In contrast, the remainder of differentially transcribed genes were only observed in evolved plasmid carrying clones, potentially representing evolved transcriptional changes driven by the plasmid (Figure 3).

**Figure 3.**
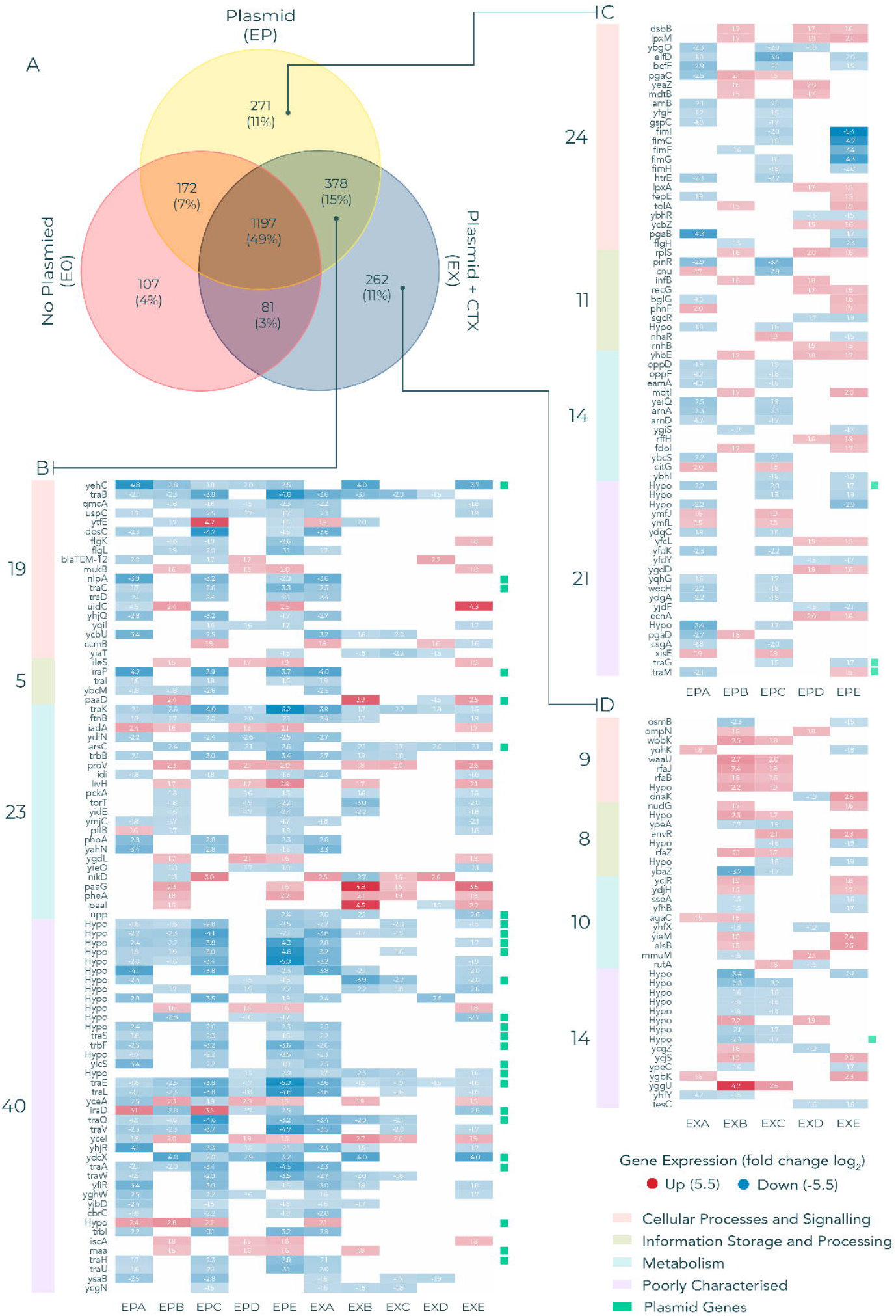
Transcriptional changes in evolved *E. coli* MG1655. Genes that were significantly differentially expressed (Log_2_ FC ≥1.5, FDR ≤0.05) in evolved clones relative to their ancestor. A) Number of significantly differentially transcribed genes in each evolution treatment. B) Expression values of significantly differentially expressed genes that are common to both the EX and EP treatments. Only genes that are present in ≥3 evolved clones per treatment are displayed to prioritise genes that were under parallel selection. C) Expression of genes that were uniquely differentially expressed in the EP treatment. D) Expression of genes that are uniquely differentially expressed in the evolved EX treatment. Red cells indicate increased expression, blue cells indicate decreased expression relative to their ancestor. Plasmid genes are demarked with green squares. A number of plasmid genes are downregulated across several replicates of both treatment conditions, including a number of genes from the *tra* operon.

Among this subset, we focused on those significant differentially transcribed in parallel in at least 3 replicate populations per treatment (n=452) to identify those most likely to represent adaptive evolved responses. In general, most of these evolved changes led to downregulation (∼80%, n=363). Several functions exhibited plasmid-associated parallel transcriptional responses, including fimbrae (*fimCFGHI yehC, yqiL*), flagellar (*flgK, flgL, yfiR*), cellular adhesion (*ycbU*), DNA damage (*ybaZ, recG, uspC*), efflux (*envR, mdtB, ybhR*), LPS (*waaU, rfaBJ, rfaZ*), outer membrane (*dsbB, ompN, htrE, lpxA, qmcA*) and biofilm (*ycgZ, pgaBC*). In addition, several plasmid genes were significantly downregulated across both treatments. Many of these were hypothetical genes, however several members of the *tra* operon were among the commonly downregulated genes, although this downregulation did not result in complete ablation of conjugational ability (Table S3).

Three plasmid-containing evolved clones exhibited a transcriptional response distinct from the others. This response was characterised by strong upregulation of Hns and strong downregulation of CTX-M-15 and was associated with insertions of IS1 or IS3 between *hns* and *tdk*. Notably, the evolved clones with these IS-element insertions and consequently downregulated CTX-M-15 expression tended to show reduced resistance to cefotaxime compared to the ancestral MG1655 (pLL35) strain (Figure 4).

**Figure 4.**
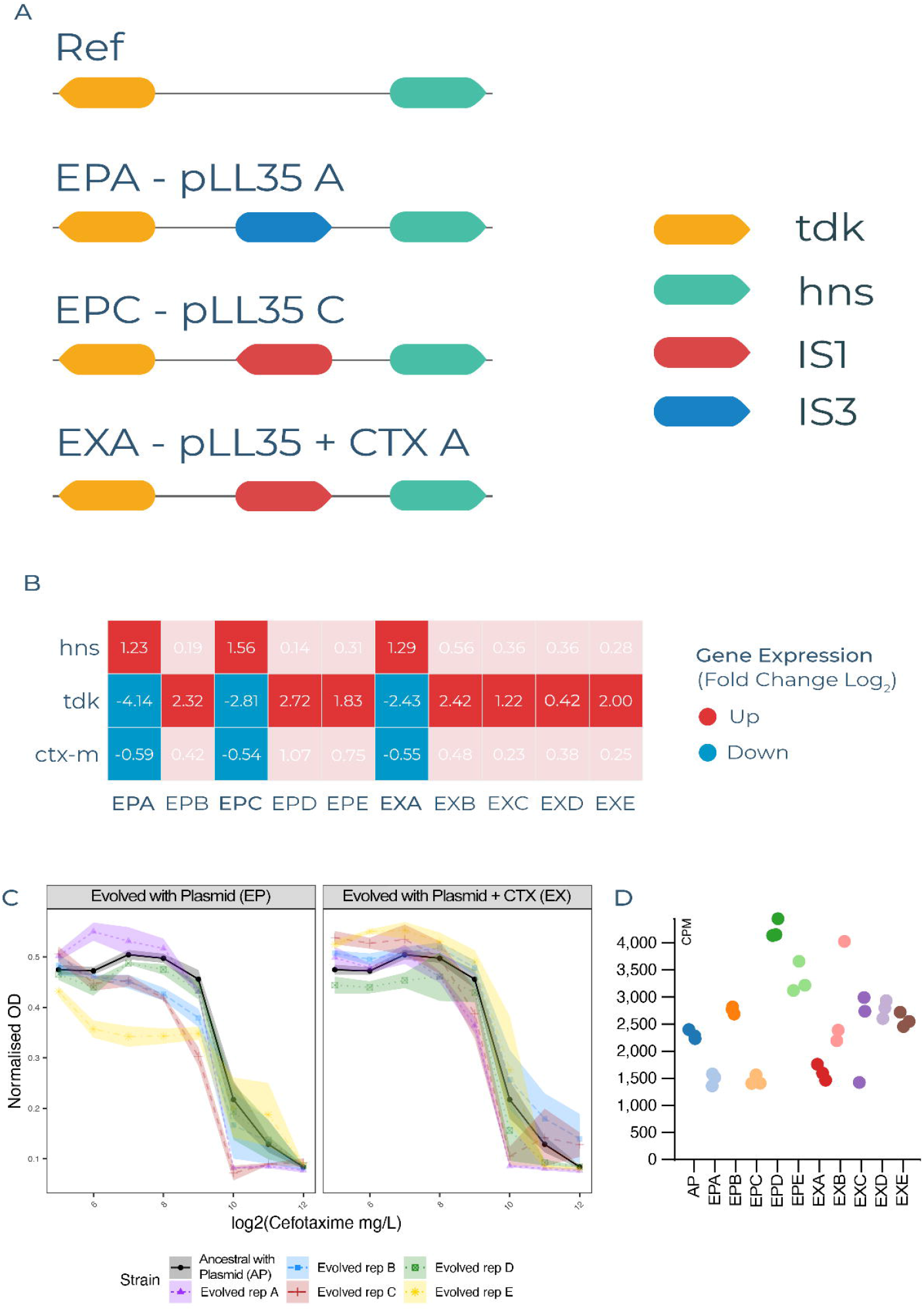
Transcriptional and resistance effects of IS insertions upstream of HNS. Orientation and location of IS-element insertions between *hns* and *tdk* in 3 *E. coli* MG1655 plasmid-carrier evolved clones. A) The reference configuration of the region, and the insertion position and orientation of IS-elements in evolved clones. B) Differential expression of *hns, tdk* and *bla*_CTX-M-15_ for all MG1655 plasmid-carrier evolved clones relative to their ancestor. Red cells indicate increased expression, blue cells indicate decreased expression relative to their ancestor. C) Minimum inhibitory concentration curves against cefotaxime showing variation between the individual evolved and ancestral clones for the EP and EX treatments (different evolving lines denoted by colours; see visual key). D) Normalised transcript counts of *bla*_CTX-M-15_ (counts per million) with the different evolving lines denoted by colours (see visual key).

## Discussion

As plasmids transmit between bacterial lineages in microbial communities, they encounter diverse genomic backgrounds, wherein a given plasmid can have different fitness effects and cause varying levels and types of cellular disruption among strains (Alonso-del Valle et al., 2021; Dunn et al., 2021). The evolutionary response to plasmid acquisition in bacteria has been studied in a range of plasmid-host systems, providing evidence of evolutionary mechanisms which explain the widespread existence of plasmids in bacterial genomes (Brockhurst and Harrison, 2022; San Millan, 2018). However relatively few of these studies have compared evolutionary responses to gaining the same plasmid between genetically divergent strains, nor done so for a multi-drug resistance plasmid in clinically relevant bacteria. In this study we examined the evolutionary response to plasmid acquisition after 700 generations by experimentally evolving genetically divergent *E. coli* lineages—including environmental, clinical and lab strains—with the MDR plasmid pLL35 in the presence and absence of antibiotic selection pressure.

We show that pLL35 acquisition imposed a small initial fitness cost in all the divergent *E. coli* strains we used. However, in only some of these strains did evolved clones show improved growth relative to their ancestor, notably strains MG1665 and F022, and such improvements were strongest in replicates evolved with cefotaxime. In addition, no consistent evolved changes to cefotaxime resistance were observed among strains, albeit with high divergence among replicates in some strains (e.g., MG1655). Nevertheless, by filtering for mutations that occurred exclusively in evolved plasmid carriers (i.e., those that did not occur in the corresponding plasmid-free control) and focusing on loci mutated in multiple replicate lines, we identified a range of chromosomal mutations putatively associated with adapting to plasmid carriage, some of which were common to multiple strains. These chromosomal mutations affected a wide range of operons but were enriched for metabolic and regulatory cellular functions. Plasmid mutations were less numerous and most commonly affected conjugation-related genes. On both the chromosome and the plasmid, we observed an important role for IS-mediated mutations in some strains, including insertion mutations impacting global regulatory systems (e.g. HNS) with effects on cefotaxime resistance unique to these evolved clones.

The majority of parallel chromosomal mutations associated with adapting to the plasmid occurred in genes involved in metabolism or its regulation. Several of these genes and operons include those that showed a transcriptional response in our previous RNAseq study of the immediate impact pLL35 acquisition in these strains, including *artP, nha*, and *put* (Dunn et al., 2021). A notable finding of our previous study was a consistent low-level upregulation of genes involved in anaerobic metabolism caused by pLL35 across all the genetically divergent strains (Dunn et al., 2021). Consistent with this conserved transcriptional response, here we observed parallel mutations in a wide range of genes associated with anaerobic metabolism, including anaerobic respiration (*norR*), fatty acid metabolism (*fad*) and fumarate metabolism (*dct*) in evolved plasmid carriers from multiple strains. This adds to a growing body of evidence suggesting that metabolism and the evolution of MDR *E. coli* are intrinsically linked. A pangenomic analysis of the MDR clone of *E. coli* ST131 showed that the MDR clone was enriched in allelic variation in core anaerobic metabolism genes (McNally et al., 2019).

Additionally, a study of diverse *E. coli* lineages grown under sub-inhibitory antibiotic selection pressure resulted in parallel adaptations in metabolism genes which potentiated resistance (Lopatkin et al., 2021), and recent modelling studies have shown that evolution of AMR is strongly coupled to the evolution of major metabolic pathways (Pinheiro et al., 2021). More generally, transcriptional and metabolomic studies suggest that a wide range of plasmids impact cellular metabolism in diverse bacterial taxa (Vial and Hommais, 2020). Why plasmids commonly alter cellular metabolism is currently unclear, but this may reflect a plasmid strategy designed to remodel bacterial metabolism in ways that are niche adaptive and boost plasmid vertical transmission (Billane et al., 2022). Our data suggest that, over the longer term, bacterial hosts may respond to plasmid metabolic manipulation through compensatory mutations in the affected metabolic pathways.

Although plasmid mutations were less commonly observed than those affecting chromosomal genes, plasmid mutations did arise in three of the *E. coli* strain backgrounds and almost all occurred in genes linked to conjugation. In one case, we observed the complete deletion of the conjugation operon, rendering the plasmid non-conjugative. Additionally, our RNAseq data showed that evolved MG1655 plasmid carriers near universally downregulated genes encoding components of the plasmid conjugation machinery. The evolution of reduced conjugation is a common mechanism by which plasmids adapt to long-term association with a given bacterial host (Dahlberg and Chao, 2003; Porse et al., 2016; Turner et al., 2014), and this is likely to reflect the burden of expressing the conjugative machinery for host cells and the absence of horizontal transmission fitness benefits for plasmids in populations without a supply of plasmid-free recipients cells (Hall et al., 2017a). Notably, loss of conjugation was one of several possible routes of compensatory evolution for p-OXA-48 plasmids in *E. coli* that have been observed to occur within patient infections (DelaFuente et al., 2022).

Other resident mobile genetic elements (MGE), notably several insertion sequences, caused parallel mutations at a range of sites affecting both the chromosome and the plasmid in evolved plasmid carriers across multiple strains. We posit two non-mutually exclusive explanations for this pattern: First, MGE mobilisation and expansion may be triggered by plasmid acquisition.

Interactions between conjugative plasmids and resident MGEs have been observed for chromosomal transposons that respond and relocate to plasmids (Hall et al., 2021, 2017b), and have been implicated in the mobilisation and spread of IS-encoded ARGs by plasmids (Che et al., 2021). Second, IS expansions may be a faster route to adaptation than point mutations. This is consistent with previous *E. coli* experimental evolution studies, including the Long Term Evolution Experiment where an appreciable fraction of early beneficial mutations were caused by IS elements in non-mutator lineages (Consuegra et al., 2021), and coevolution of *E. coli* with MDR plasmids where adaptive mutations have been associated with IS element disruptions of chromosomal and plasmid genes (Bottery et al., 2019, 2017; Porse et al., 2016).

The mobilization of IS1 and IS3 into the region upstream of *hns* occurred in parallel lines of MG1655 resulting in upregulation of *hns* and consequent down-regulation of the CTX-M-15 gene on pLL35 and a reduction in phenotypic resistance to cefotaxime. H-NS has been shown to play a key role in the controlled acquisition and integration of a number of virulence encoding MGEs such as Salmonella Pathogenicity Islands (Ali et al., 2014) and the Locus of Enterocyte Effacement in *E. coli* O157 (Sperandio et al., 1999), with H-NS silencing transcription of operons on those MGEs to minimize their impact on fitness when acquired. To our knowledge, a role for H-NS silencing during MDR plasmid acquisition has not previously been shown in *E. coli*. Our data suggest H-NS may offer a regulatory route by which *E. coli* adapts to MDR plasmids, occurring here via the insertion of an IS element into the regulatory region of *hns* upregulating H-NS and in turn down regulating the newly acquired plasmid.

Using a comparative experimental evolution approach our study reveals a range of evolutionary pathways taken by genetically divergent *E. coli* lineages following acquisition of an MDR plasmid. In contrast to some studies of compensatory evolution to ameliorate plasmid fitness costs, we did not see evidence for a major genetic conflict between chromosome and plasmid that could be fixed by single point mutations (cf. (Hall et al., 2021)). Rather we observed a more complex evolutionary process targeting a wide range of functions, including cellular metabolic pathways impacted by plasmid carriage, plasmid conjugation, and global regulatory systems controlling expression of foreign DNA. Our results highlight the importance of interactions between incoming plasmids and MGEs already resident in the cell, both in terms of IS elements relocating to plasmids and their expansion causing parallel mutations, an emerging theme in plasmid-host evolution (Bottery et al., 2019, 2017; Porse et al., 2016). The key challenge for future work will be to understand how these evolutionary processes translate from the lab into more infection-relevant conditions to better understand the success of specific plasmid-host combinations in clinical settings.

## Materials and methods

### Bacterial strains and plasmids

Five genetically diverse *E. coli* strains were used as hosts for the multidrug resistance plasmid pLL35 (Dunn et al., 2021). Specifically, these included clinical isolates (F022, sequence type ST-131 A; F054, sequence type ST-131 B; F104, sequence type ST-131 C) an environmental isolate (ELU39, sequence type ST-1122) and the lab strain MG1655. Five independent colonies were isolated per strain for use as ancestral genotypes in the evolution experiments and stored cryogenically for subsequent use (plasmid-free ancestrals). For use in plasmid-containing treatments, pLL35 was conjugated into each of these ancestral genotypes from its natural host (*Klebsiella pneumoniae*), generating five independent transconjugants per strain (plasmid-carrying ancestrals). Specifically, each single colony was inoculated into 5 ml of nutrient broth (NB; Oxoid, United Kingdom) and incubated at 37°C for 2 h with shaking (180 rpm). *K. pneumoniae* donor and *E. coli* recipient cultures were mixed at a ratio of 1:3, and 50 μl was used to inoculate 6 ml of brain heart infusion (BHI) broth. These were then incubated as static cultures at 37°C for 24 h. The conjugation mixture was plated onto 4 μg/ml of cefotaxime UTI Chromagar (Sigma-Aldrich, United Kingdom) and incubated at 37°C overnight. Pink *E. coli* transconjugant colonies were then subcultured onto UTI Chromagar with 4 μg/ml of cefotaxime and stored cryogenically for subsequent use.

### Selection experiment

We isolated five independent plasmid-free and plasmid-carrying ancestral clones per strain. Each of these was re-streaked onto a nutrient agar plate, incubated at 37°C for 24 h, and a single colony from each plate was then used to inoculate an overnight liquid culture in 6 ml of NB. To establish replicate populations for the selection experiment, 60 μl of the respective overnight culture was used per treatment. Plasmid-carriers were propagated by 1% daily serial transfer in 6 ml NB liquid cultures under two treatments: Specifically, replicate plasmid-carrier populations were propagated either with (evolved with plasmid plus cefotaxime treatment; EX) or without (evolved with plasmid treatment; EP) 4 μg/ml of cefotaxime supplementation. In addition, plasmid-free controls were propagated under equivalent conditions without cefotaxime supplementation (control treatment; C). This experimental design resulted in 75 independently evolving lines that were maintained for 84 days. For all treatments, every 14 days serial dilutions of each population were plated out onto nutrient agar plates ± 4 μg/ml of cefotaxime to quantify the total population density and the density of cefotaxime resistance. For the EX and EP treatments, every 28 days 24 colonies from the cefotaxime supplemented agar plates were picked and tested for the presence of the plasmid and the CTX-M-15 gene by PCR using a previously published protocol (Dunn et al., 2021).

### Growth kinetics assays

Growth kinetics were obtained for each plasmid-free and plasmid-carrying ancestral clone and for a single colony randomly chosen from each evolving line at the end point of the selection experiment (i.e., day 84). Triplicate cultures of each clone were grown in 200 μl NB per well in 96 well plates incubated at 37 °C for 24 h. Optical density (OD) at 600nm was recorded every 30 minutes for 24 hours using an automated absorbance plate reader (Tecan Spark 10). Prior to each reading, plates were shaken for 5s at 180rpm orbital shaking with a movement amplitude of 3 mm. A humidity cassette was used to minimize evaporation.

### Cefotaxime resistance assays

MIC assays for each plasmid-free and plasmid-carrying ancestral clone and for a single colony randomly chosen from each evolving line at the end point of the selection experiment (i.e., day 84) were conducted according to the CLSI guidelines (Wikler et al., 2006), using nutrient broth and cefotaxime. Shaking (180 rpm) overnight cultures in 6 ml nutrient broth were established from independent colonies previously grown on agar plates. The following day, 0.5 McFarland cell suspensions were prepared and further diluted 1/500 to inoculate 200 μl of nutrient broth in 96-well plates. The final cefotaxime concentrations tested were 2-fold increases from 64 to 8192 μg/ml, and the final volume per well was 200 μl (100 μl bacterial inoculum plus 100 μl antibiotic solution). The OD_600_ was recorded after 24 h of static incubation at 37°C and normalized by subtracting the OD_600_ of a blank well. As working with positive OD_600_ values facilitates further data analysis and interpretation, we linearly transformed OD_600_ estimates by adding 0.0819 to all data. For each strain and plasmid combination, the relative growth at each antibiotic concentration was obtained by dividing OD_600_values in the presence of antibiotic by the OD_600_ of the corresponding parental strain grown in the absence of antibiotic. The relative growth values were used to calculate the area under the curve (AUC) with the auc function from the R package flux (flux_0.3-0). Statistical analyses were performed on Box Cox transformed data to fulfil ANOVA assumptions.

### Statistical Analysis

RStudio was used to perform statistical analysis. The bacterial densities from the evolution experiment were analysed in a linear mixed-effects model (LMM) with the R package “nlme”. Strain, treatment and transfer were introduced in the model as fixed effects, whereas population was used as a random effect to account for the repeated samplings. Data required a Box-Cox transformation to meet model assumptions. We reduced the model by performing likelihood ratio tests on nested models. We found a significant interaction effect between strain and transfer.

The impact of strain and evolution treatment on growth kinetic parameters including the area under the curve, the maximum growth rate, the maximum optical density and the lag time of the evolved strains relative to their corresponding ancestral were assessed. These relative growth parameters were analysed using ANOVA, art ANOVA or Scheirer-Ray-Hare test depending on the data characteristic and if the assumptions of the tests were met. A Box-Cox transformation was applied when required. The impact of treatment for each of strains was also studied due to the impact that the strain had on the growth parameters.

For MIC analysis, an art ANOVA for MIC analysis regarding the area under the curve and an ANOVA for resistance fold change were performed to study the impact of strain and treatment during the experimental evolution in relation to the corresponding ancestral. Further statistical analyses were performed for each strain individually.

### Genomics and bioinformatics

We obtained whole genome sequences for one randomly chosen colony per evolving line taken from the end point of the selection experiment (i.e., day 84) using both Illumina and Oxford Nanopore Technology platforms. Sequencing data is available under BioProject accession number PRJNA848631, and a list of isolate accession numbers is provided in Table S4.

Ancestral WGS data are available under Bioproject Accession number PRJNA667580, and are described in (Dunn et al., 2021).

The long reads were processed with FiltLong (V 0.2.0) to remove short or low quality reads (<1000 bp, <Q6), and chimeric reads were removed using Unicycler’s (V 0.4.7) scrub module. The distributions of qualities and lengths of reads were assessed and visualised using NanoPlot (V 1.20.0). Where there was an abundance of reads, the data was subsampled to 100X coverage to reduce computational cost. Assemblies were constructed using Unicycler (V 0.4.7) using normal bridging mode. In a minor number of instances, small repetitive elements were not fully resolved, to address this, we also assembled our isolates with Trycycler (V 0.4.1), which uses iterative subsampling and assembly of differential read sets, and obtains a consensus. These assemblies were then polished using Racon (V1.4.10), Medaka (V 1.0.3), and Pilon (V 1.23) (implemented through unicycler_polish), using both long and short reads respectively. The genomes were annotated using Prokka (V 1.14.6). Assemblies were screened for insertion sequences using isescan (V 1.7.2), which identified a number of novel IS movements. These movements were further investigated using Artemis Comparison Tool, and adjacent sections of sequence were identified using both the annotation files and BLAST against the non-redundant bacterial protein database.

Variants were called against fully resolved and annotated assemblies using Breseq (V 0.35.4), using additional flags to ensure a minimum variant base depth of 10, quality of 20, and allele frequency of 0.9. We also assessed structural variation using Sniffles (V 1.0.12) and Assemblytics (commit #58fb525). Any deletions were verified by mapping the Illumina read sets against the assembly using Bowtie2 (V 4.8.2) and identifying regions of missing coverage. For our final list of variants, we masked any variants that occurred in the evolved no plasmid (E0) lines, as these represent basal adaptation to the media and laboratory conditions. Protein annotations in which a variant was detected were confirmed using BLAST, and where possible hypothetical proteins were assigned putative functions or families. Data visualisation as conducted using R (V 3.4.3) and ggplot (V 3.3.3), and GraphPad Prism (V 7).

### Transcriptomics and bioinfomatics

We obtained transcriptomes for MG1655 plasmid-free and plasmid-carrying ancestral clones and for each sequenced MG1655 evolved clone taken from the end point of the selection experiment (i.e., day 84). Triplicate shaken cultures were grown at 37°C to an OD600 of 0.6 in 10 ml of nutrient broth and centrifuged. Residual media was discarded, and the cell pellet was snap-frozen. Samples were shipped to GeneWiz, who performed the RNA extractions, and sequenced the purified RNA on an Illumina NovaSeq configured to 2 × 150bp cycles.

Kallisto (V 0.46.0) was used to quantify differential gene expression, with the high-quality hybrid *de novo* assemblies of the relevant ancestral strain was used as a reference. Input files were prepared using Prokka (V 1.13.3) for annotation, genbank_to_kallisto.py (https://github.com/AnnaSyme/genbank_to_kallisto.py) to convert the annotation files for use with Kallisto, and GNU-Parallel (V 20180922) for job parallelization. Differential gene expression was analyzed using Voom/Limma in Degust (V 3.20), with further processing of the resulting differential counts in R (V 3.5.3). Functional categories (COG, GO-terms) were assigned to genes using eggnog-mapper (V 2).

## Supporting information

Experimental data

## Open access statement

For the purpose of open access, the author has applied a Creative Commons Attribution (CC BY) licence to any Author Accepted Manuscript version arising.

## Data Access Statement

All experimental data sets are provided in the Supplementary Information of this article. Sequencing data is available under BioProject accession number PRJNA848631, and a list of isolate accession numbers is provided in Table S4. Ancestral WGS data are available under Bioproject Accession number PRJNA667580, and are described in (Dunn et al., 2021).

## Figure legends

**Figure S1.**
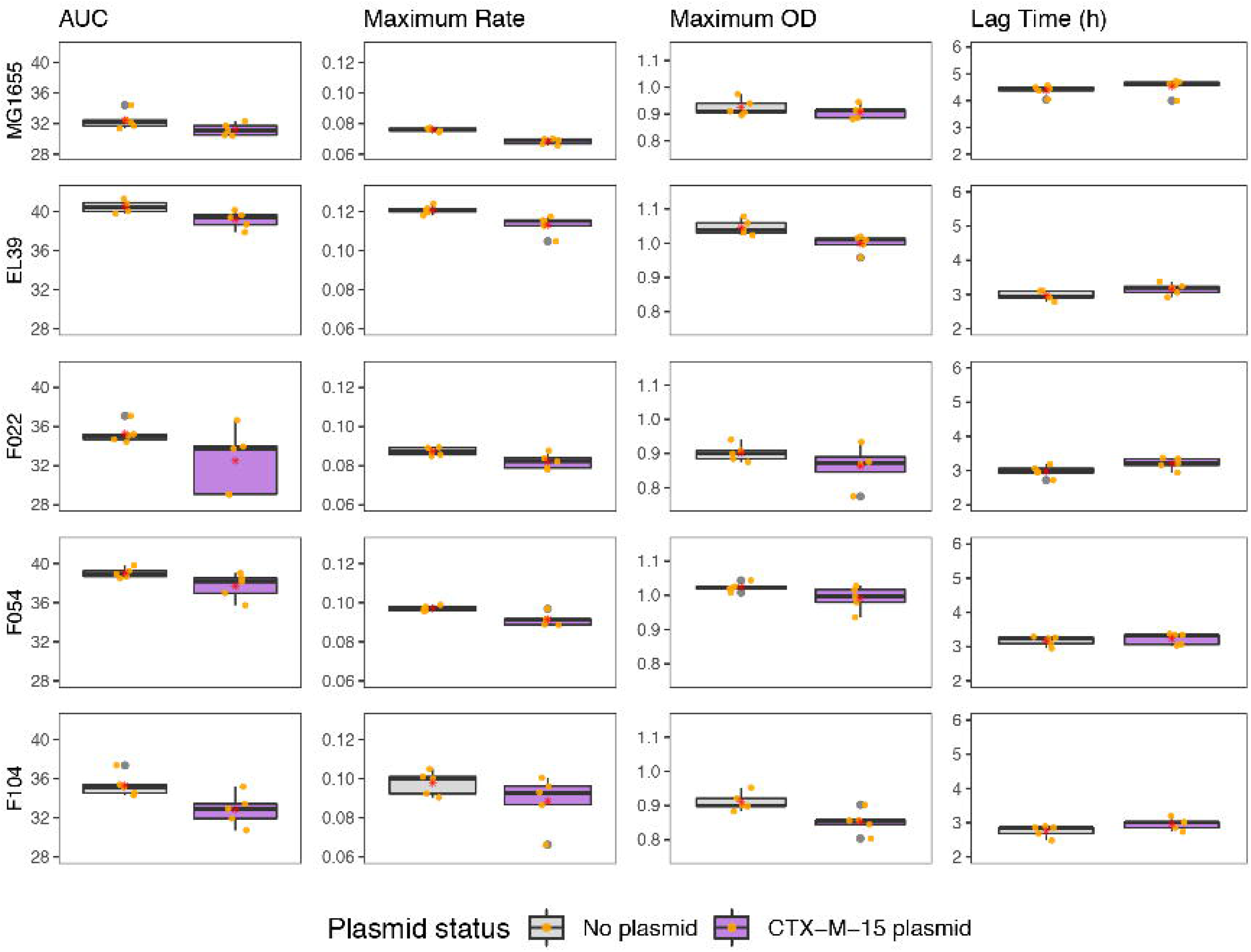
Effects of pLL35 acquisition on the growth kinetics of ancestral strains. Boxplots represent the growth kinetic parameters for the ancestral plasmid-free (grey) and ancestral plasmid-carrier (purple) clones per strain. Each strain is shown in a separate row and each growth kinetic parameter is shown in a different column, as indicated by the labels. Datapoints show the mean of technical replicates for each individual evolved clone.

**Figure S2.**
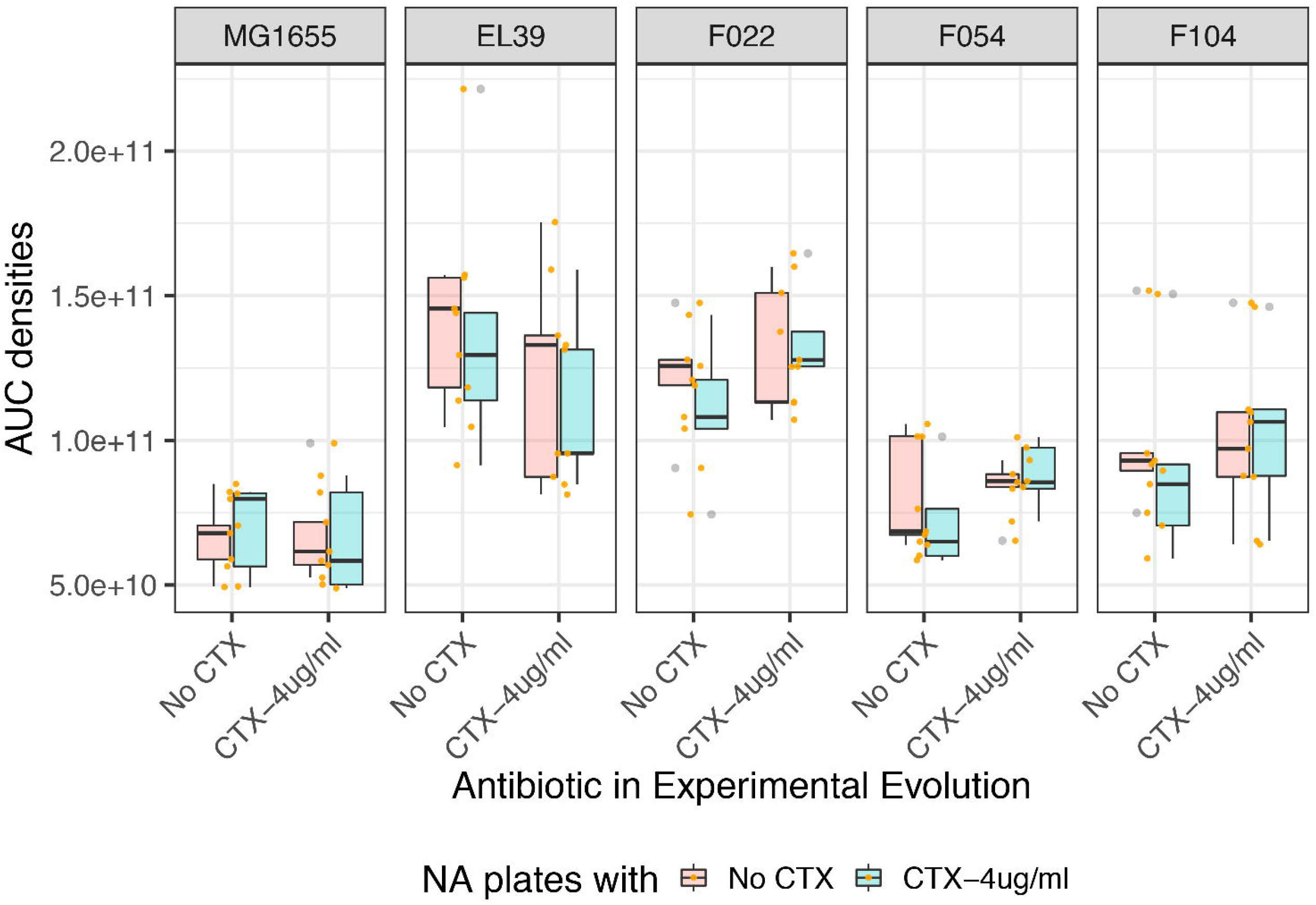
Total and cefotaxime resistant population densities. Boxplots show cumulative colony forming unit counts from nutrient agar plates (red) indicating total bacterial population density or nutrient agar plates supplemented with cefotaxime (blue) indicating the cefotaxime resistant bacterial population density calculated as area under the curve of densities over time. Strains are shown in separate panels as indicated by labels.

**Figure S3.**
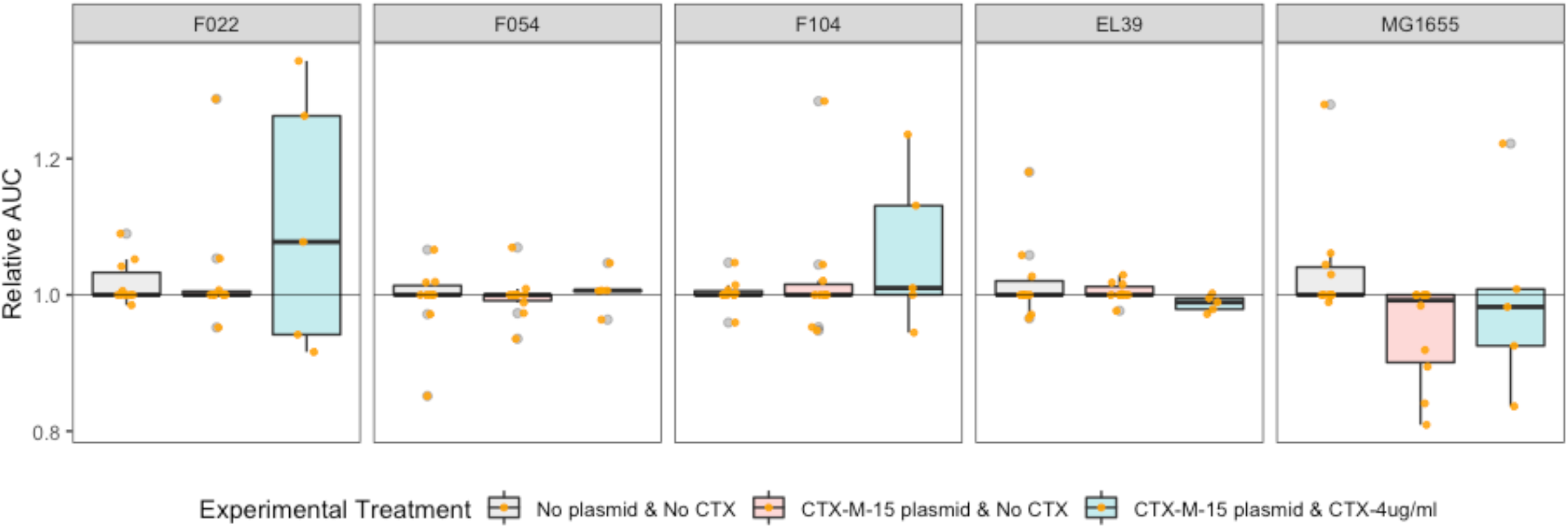
Cefotaxime resistance of evolved bacterial clones relative to their ancestor. Boxplots represent the change in cefotaxime resistance following evolution as area under the minimum inhibitory concentration curve for evolved clones relative to their ancestor. Each strain is shown in a separate panel as indicated by the label. Each evolution treatment is denoted by a colour (grey, plasmid-free control, C; red, plasmid-carrier without cefotaxime, EP; blue plasmid-carrier with cefotaxime, EX). Datapoints show the mean of technical replicates for each individual evolved clone.

**Figure S4.**
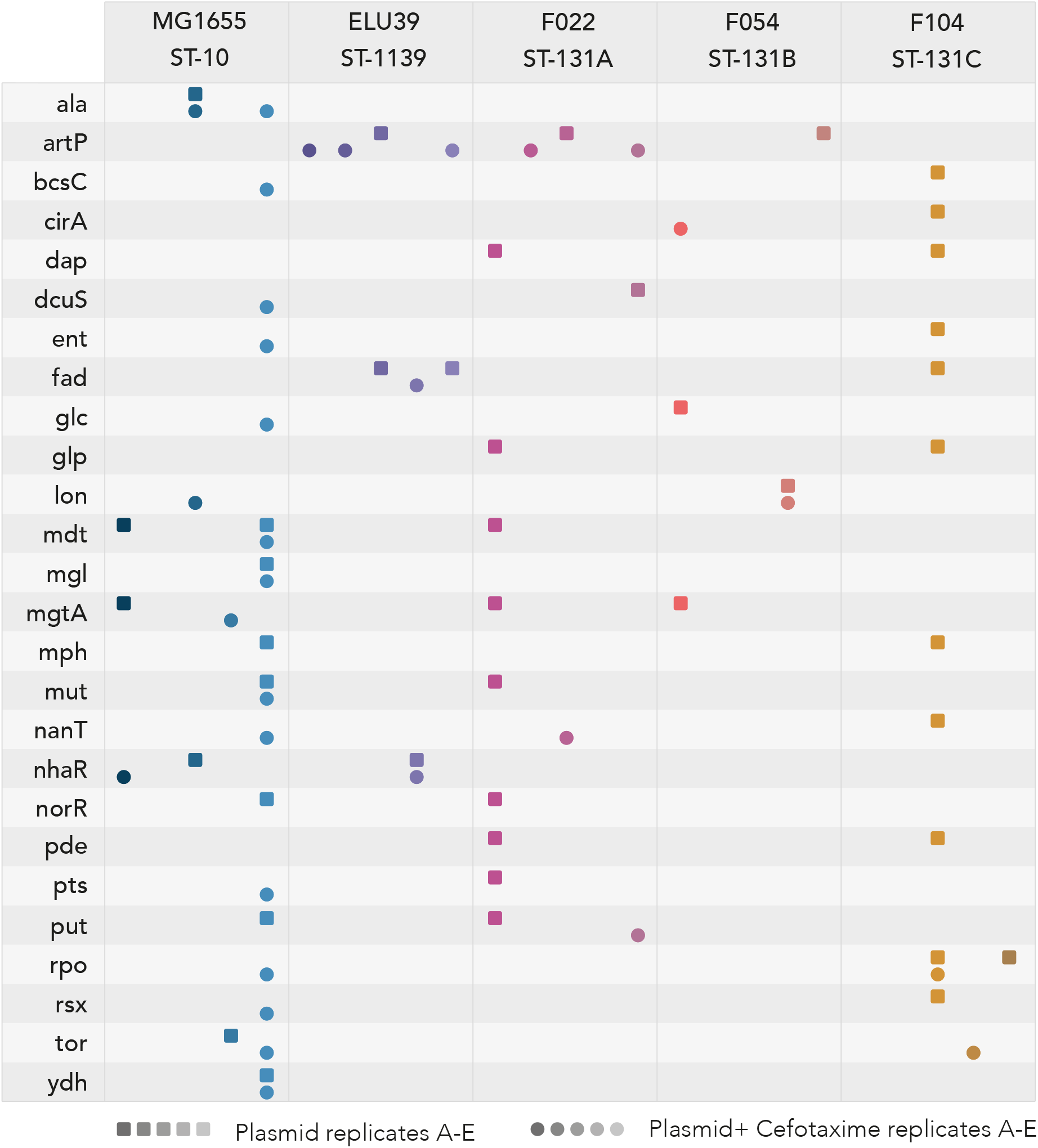
Loci and operons evolving in parallel among plasmid-carriers across strains. Chromosomal operons or genes in which mutations were detected in at least 2 independent evolved plasmid carrier clones but not in the corresponding evolved plasmid-free control. In instances where more than one gene within a given operon contained variants, the operon is listed. Where no other genes within an operon contain variants, the individual gene is listed, identifiable by the capitalised suffix.

**Figure S5.**
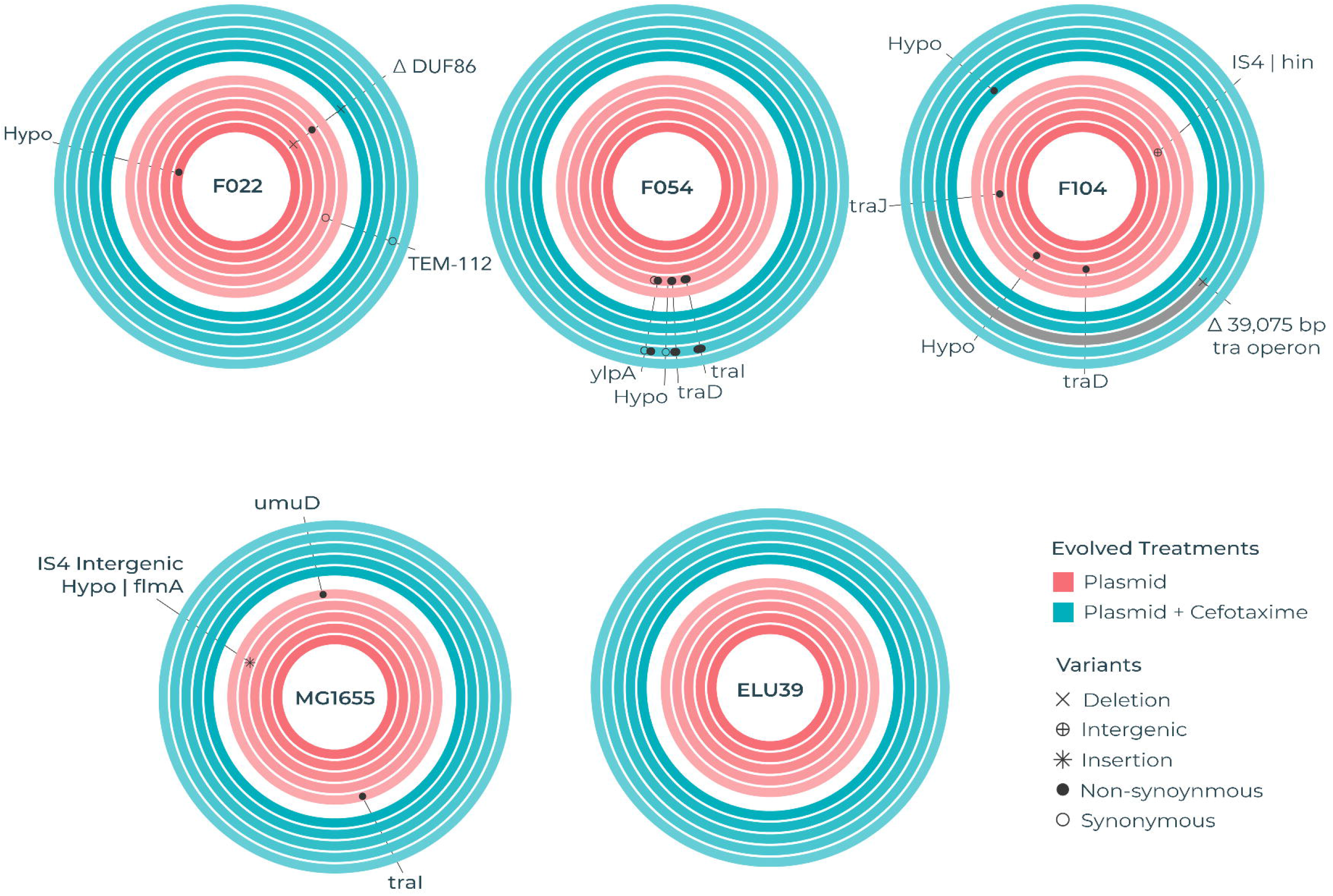
Plasmid loci targeted by parallel mutations in evolved plasmid-carriers. Each circular track represents the plasmid of an independently evolved clone. A single randomly chosen clone was genome sequenced per evolving line. As such the five replicate evolved clones per strain (as labelled) per treatment (Blue for plasmid-carriers evolved with cefotaxime, EX; Red for plasmid-carriers evolved without cefotaxime, EP) are shown as concentric tracks. Loci that acquired mutations during evolution in plasmid-carriers but not in the corresponding plasmid-free control are shown by markers denoting the type of mutation (see visual key). Loci that acquired parallel mutations in multiple independently evolving lines per strain per treatment have been labelled with the corresponding gene name or locus tag.

**Figure S6.**
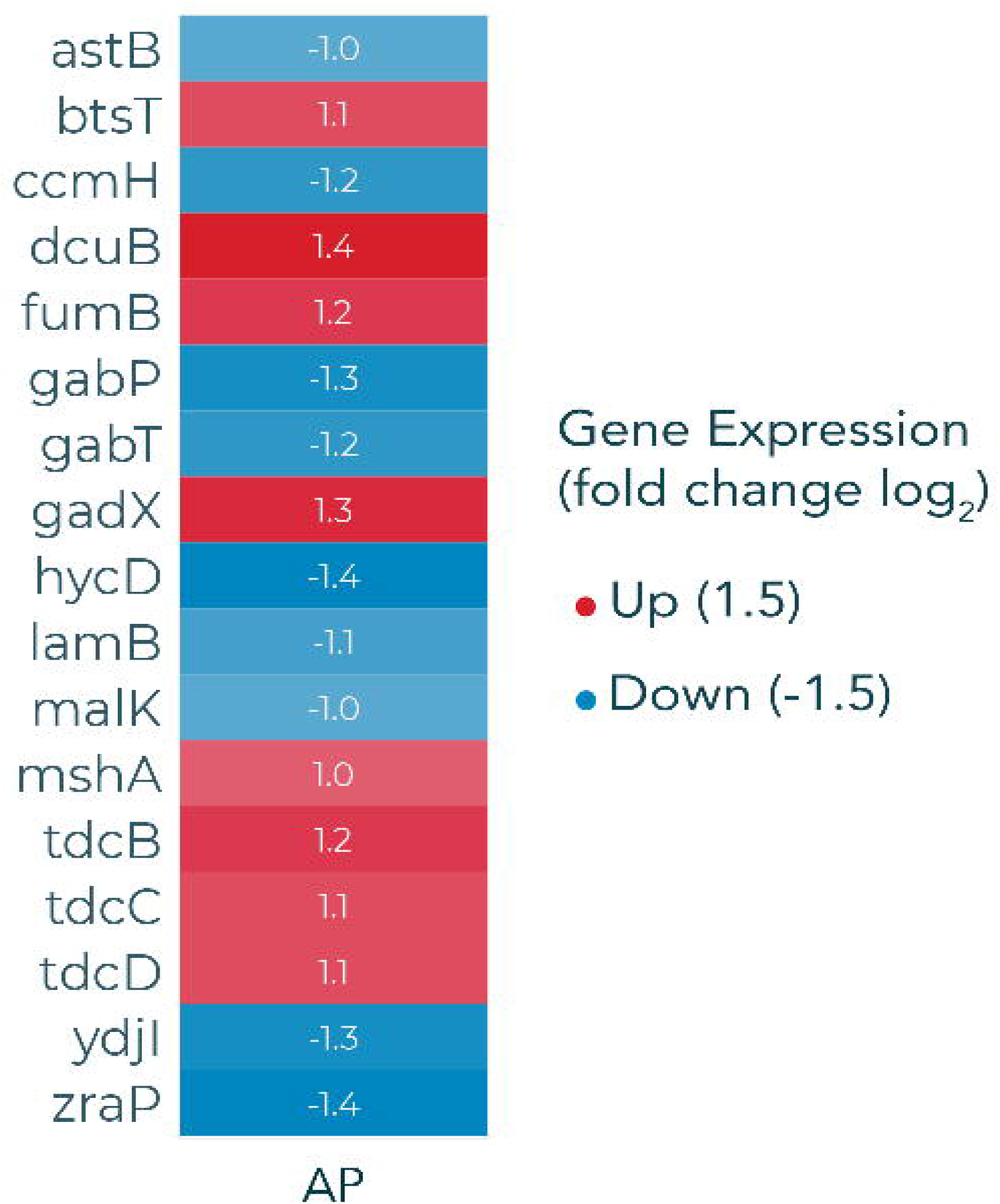
Transcriptional effect of pLL35 acquisition in ancestral *E. coli* MG1655. Genes that were significantly differentially expressed (Log_2_FC ≥1.5, FDR ≤0.05) in ancestral plasmid carrier relative to ancestral plasmid-free *E. coli* MG1655. Red cells indicate increased expression, blue cells indicate decreased expression relative to their ancestor.

**Table S1.**
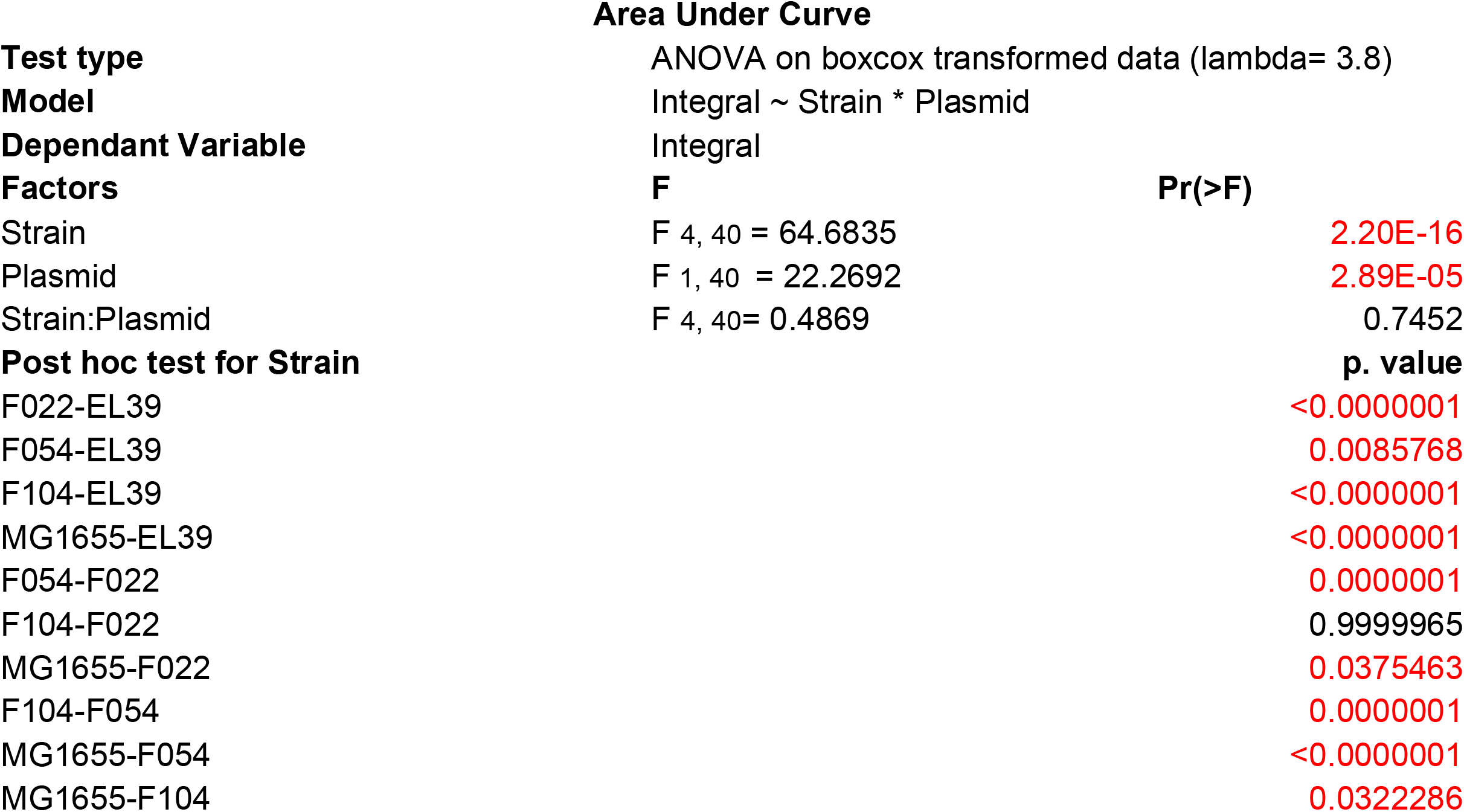
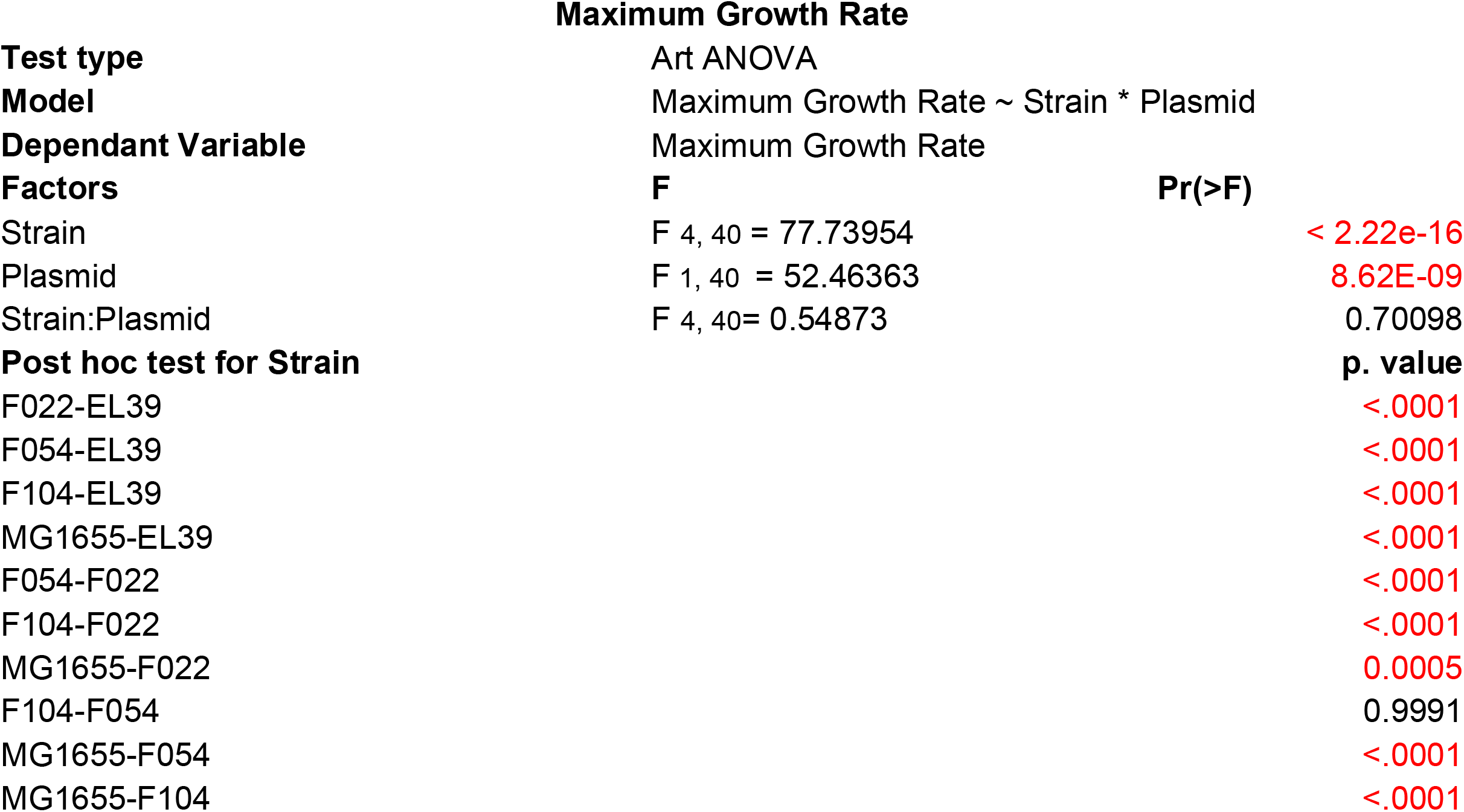
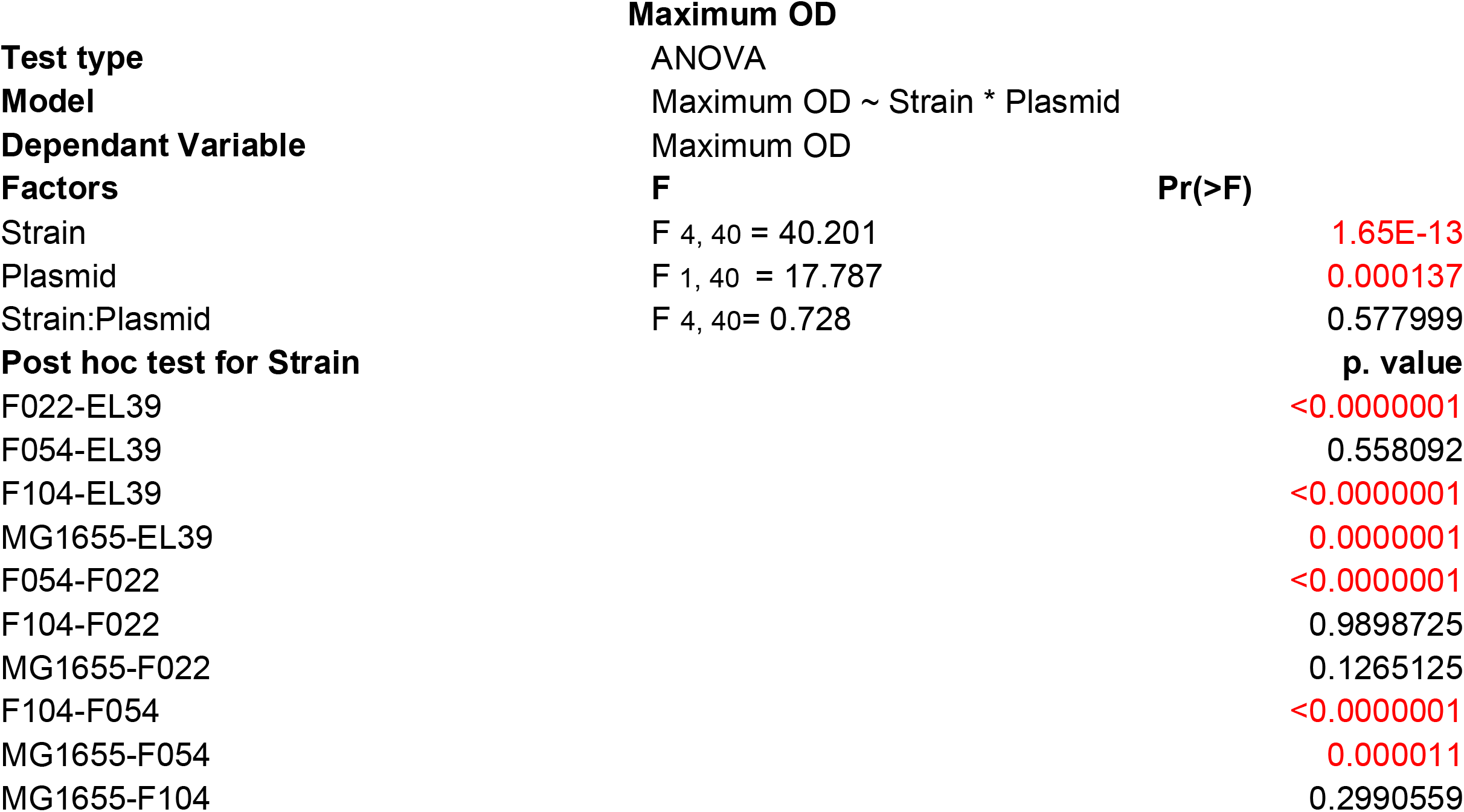
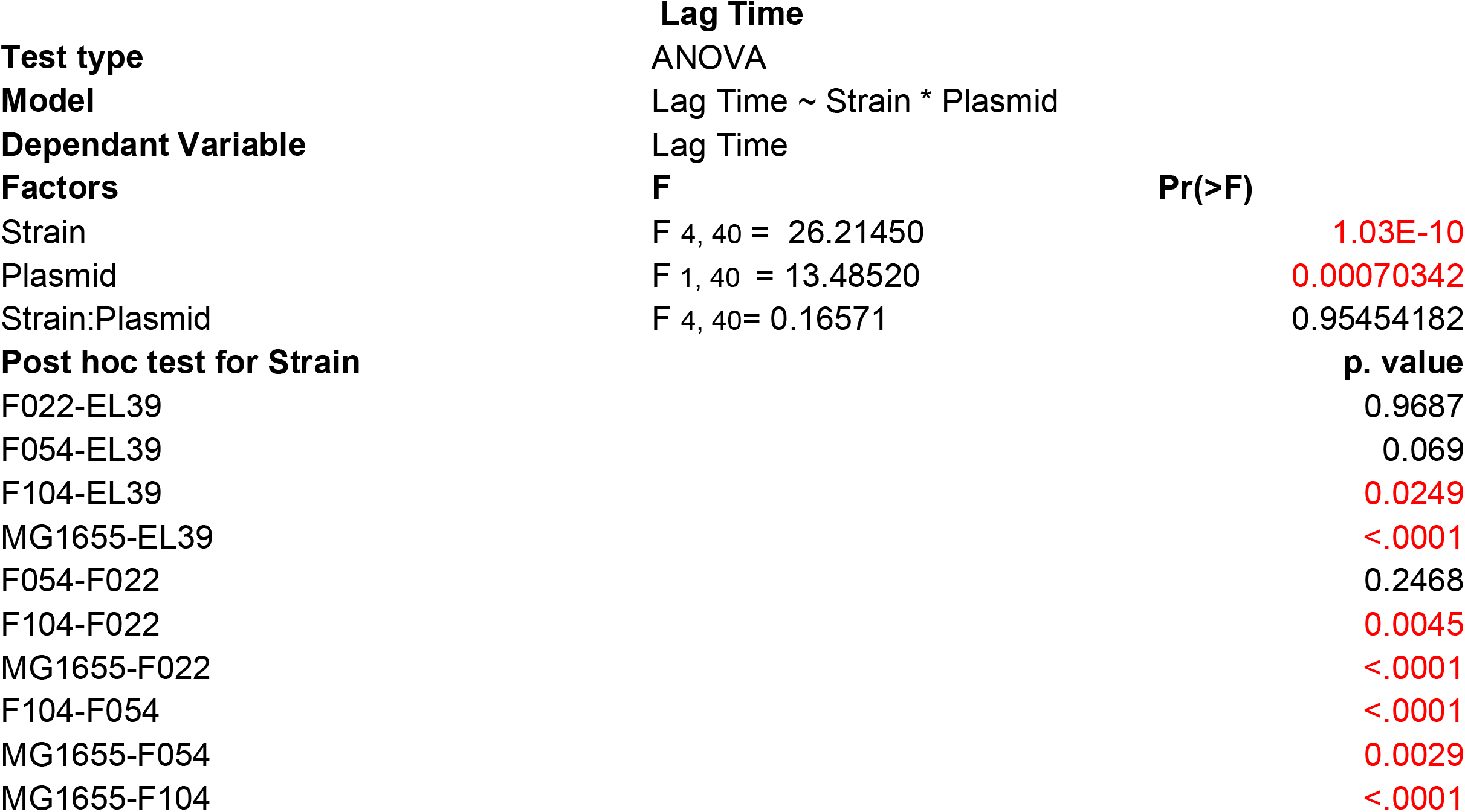
Statistical tables for ancestral growth kinetic parameters.

**Table S2.**
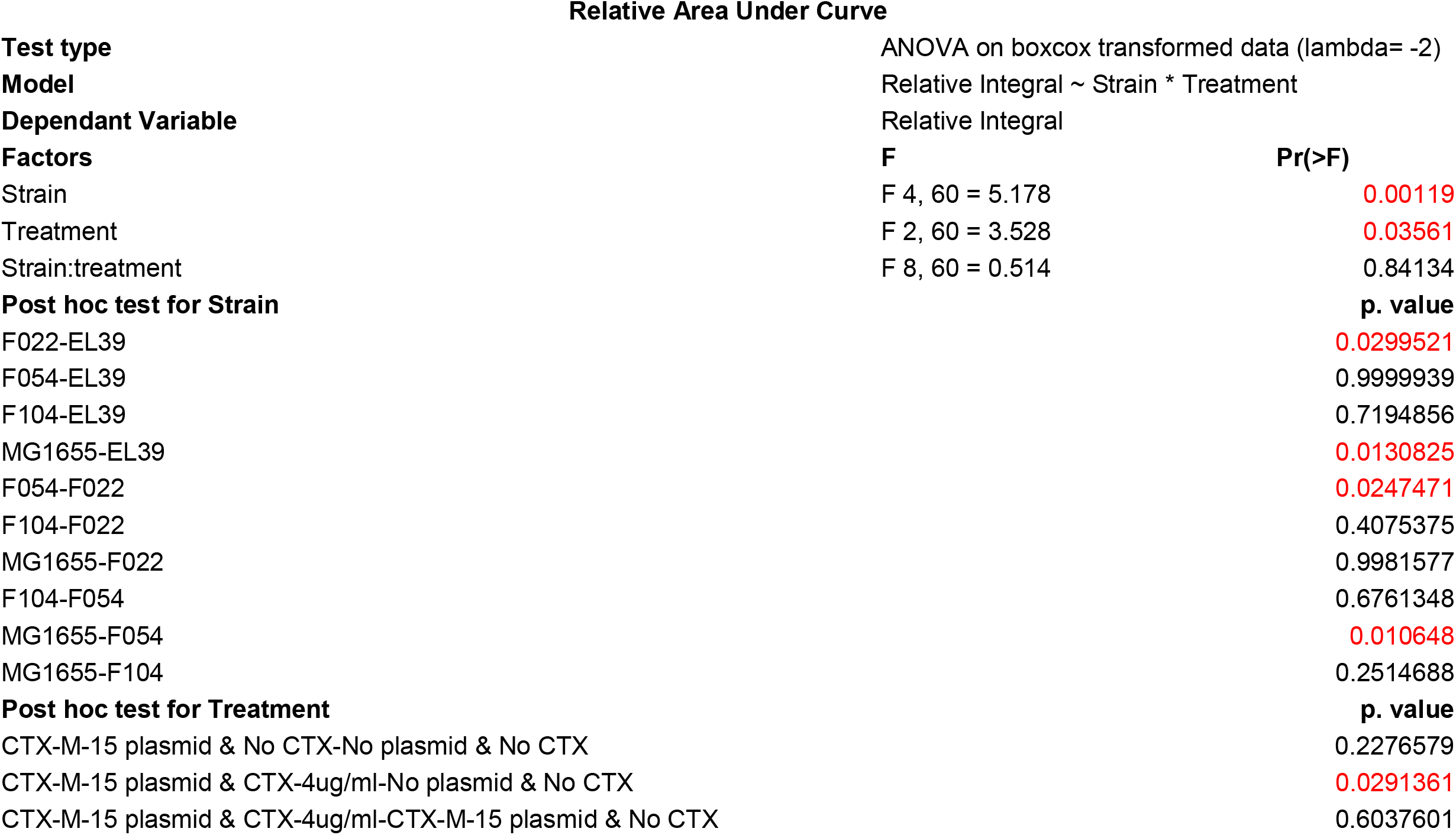
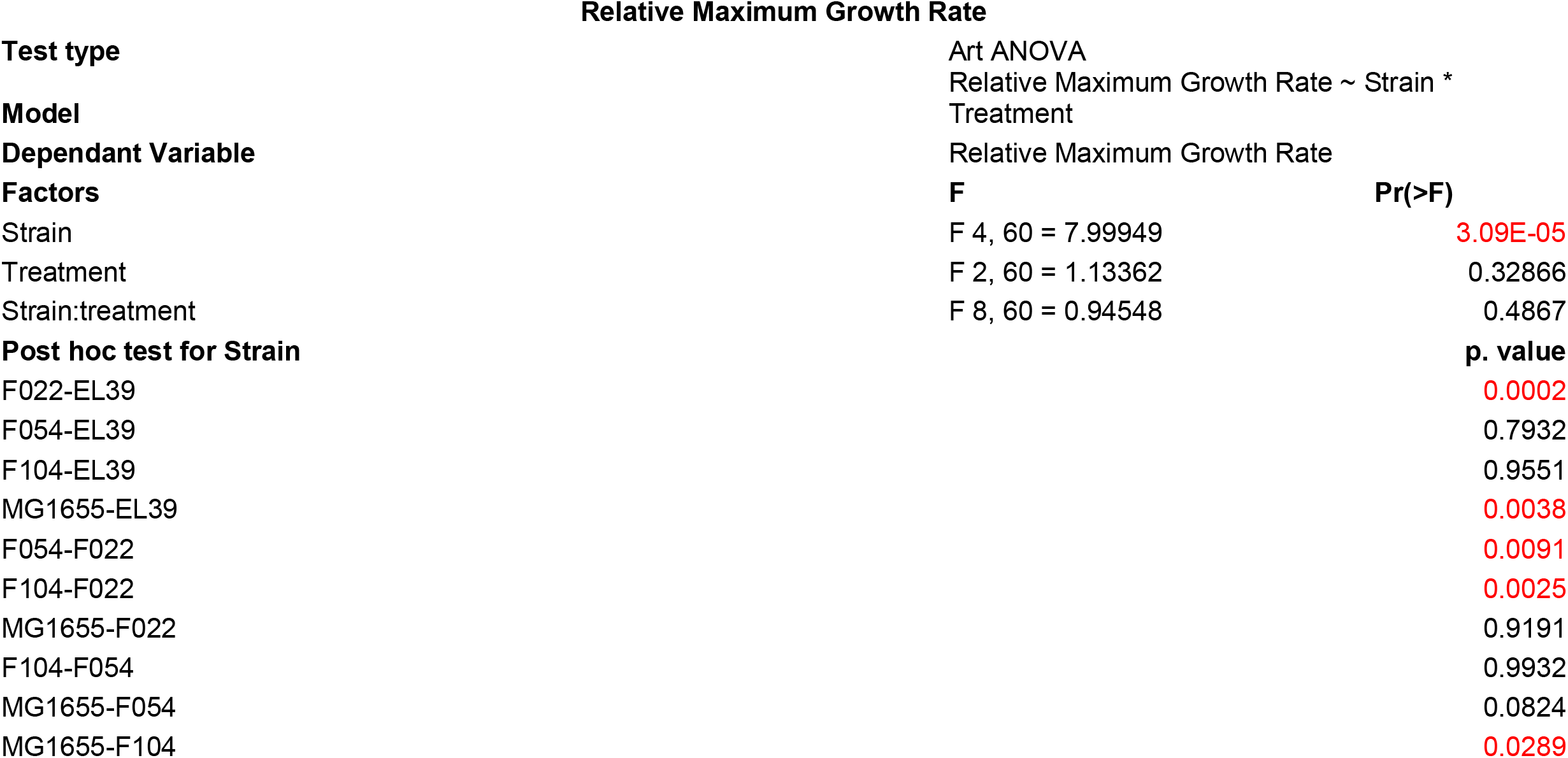
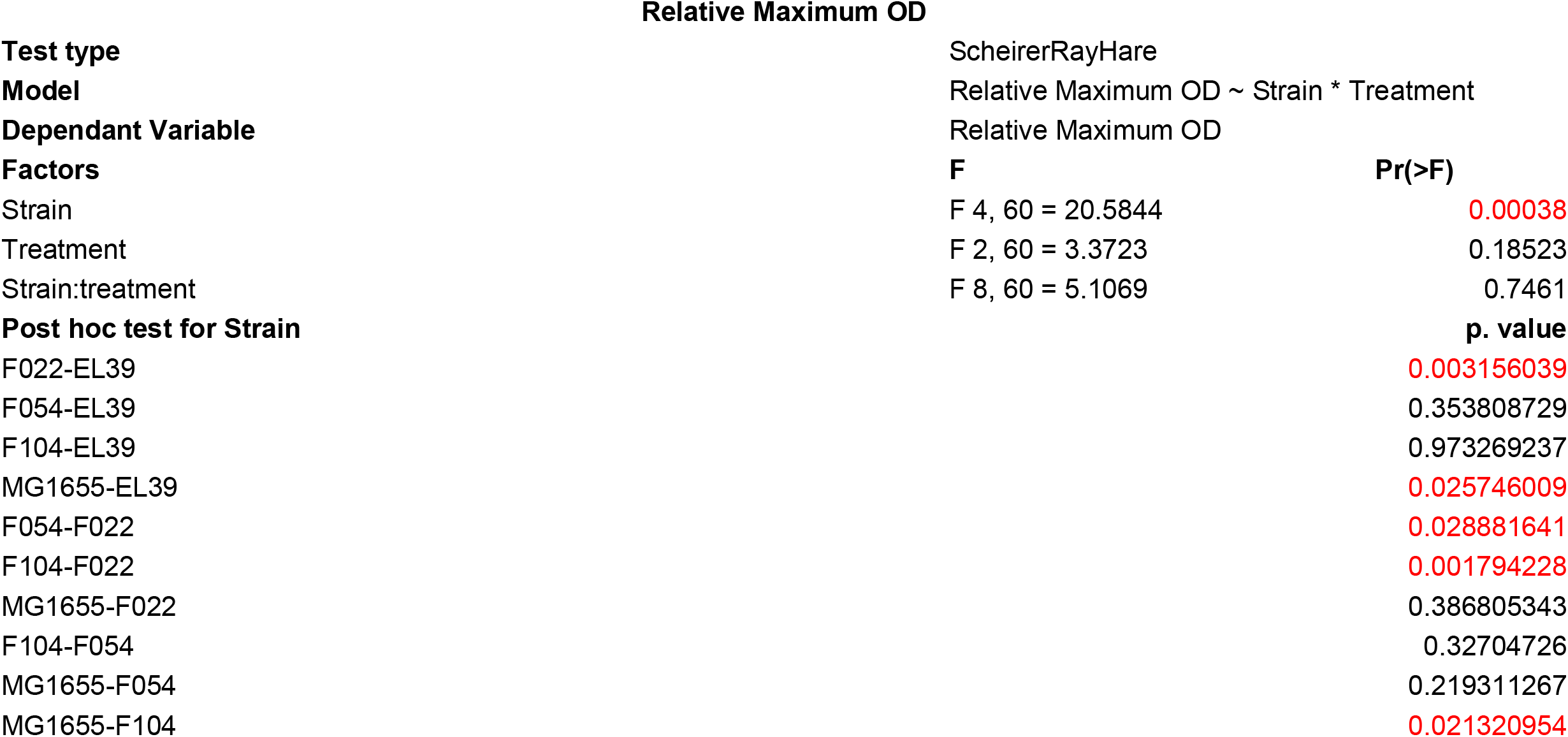
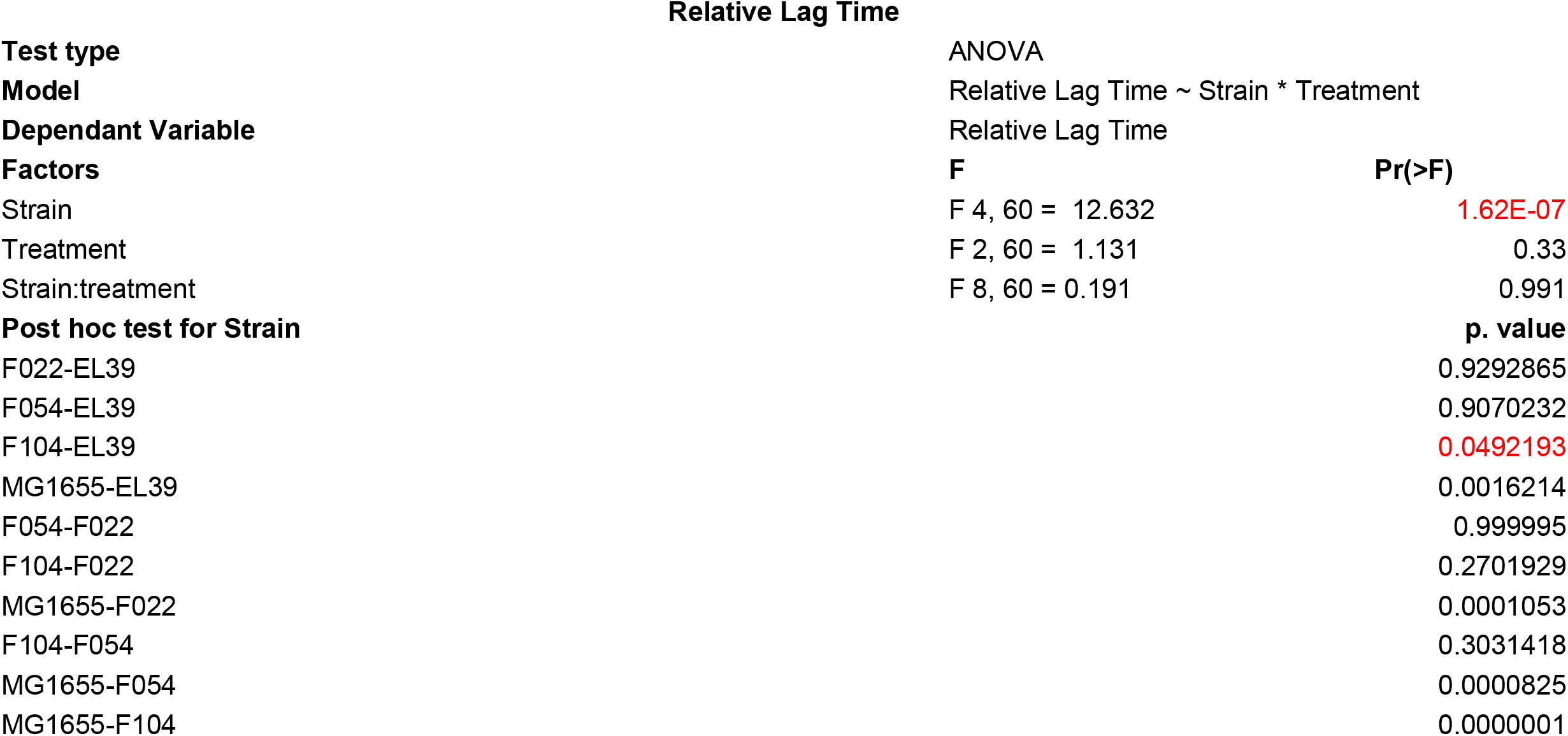
Statistical tables for e olved growth kinetic parameters.

**Table S3.**
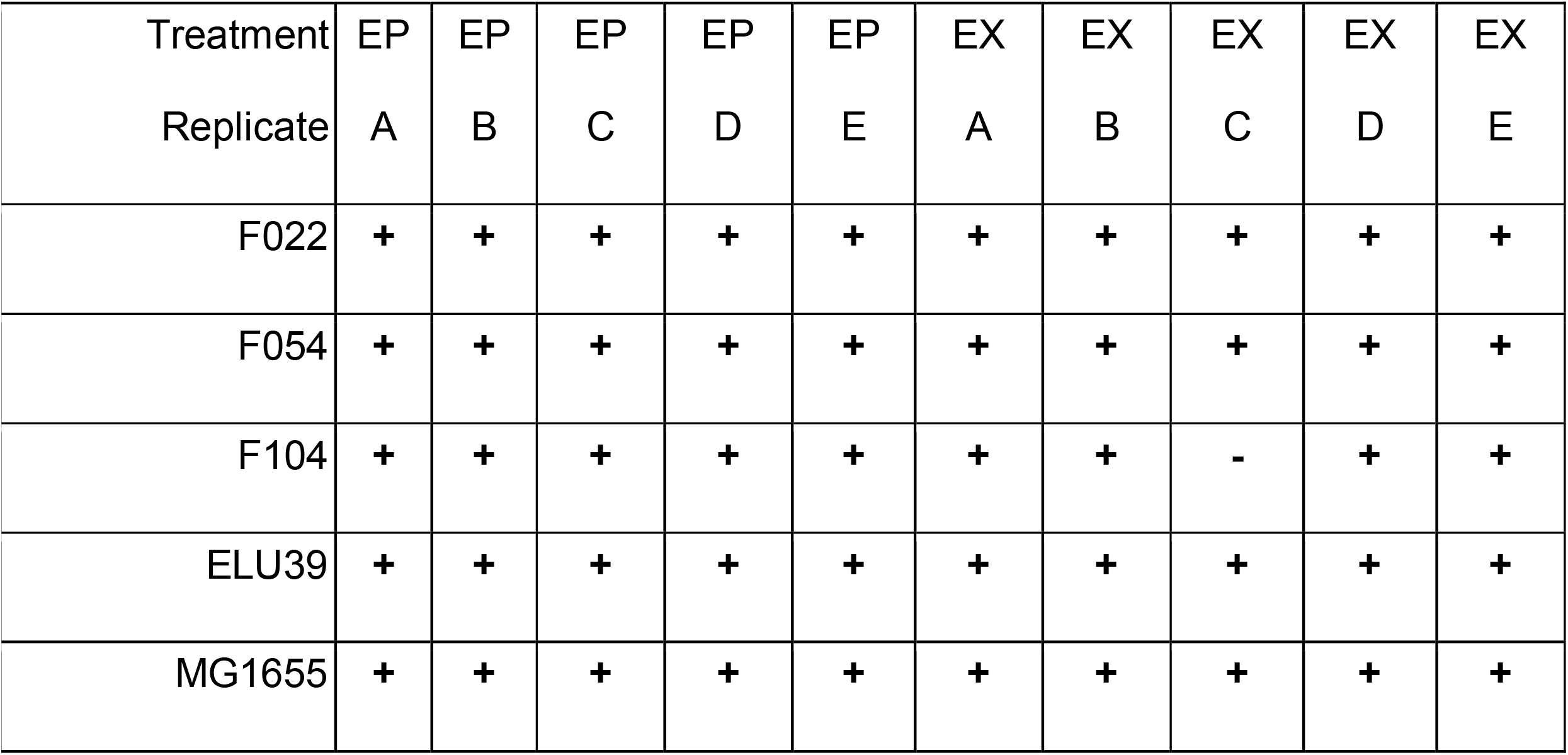
Conjugational abilities of evolved clones. Evolved plasmid-carrier clones were tested for their ability to conjugate pLL35 into an *E. coli* J53 recipient strain as previously described (Dunn et al. 2021). Conjugation positive strains are demoted with “+” and conjugation deficient strains are denoted with “−”.

**Table S4 | Short Read Archive accession numbers**

**Supplementary Table 1.** Whole genome sequencing data generated for this work is available under Bioproject Accession number PRJNA848631. Isolate suffixes relate to treatment condition; E0 = Evolved without plasmid, EP = Evolved with plasmid, EX = Evolved with plasmid in the presence of cefotaxime. Ancestral WGS data are available under Bioproject Accession number PRJNA667580, and are described in Dunn et al., 2021 (https://doi.org/10.1128/mSystems.00083-21).

| Isolate | Biosample | Illumina | Nanopore |
| --- | --- | --- | --- |
| F022_E0A | SAMN29006941 | SRR19633518 | SRR19634252 |
| F022_E0B | SAMN29006942 | SRR19633517 | SRR19634251 |
| F022_E0C | SAMN29006943 | SRR19633506 | SRR19634240 |
| F022_E0D | SAMN29006944 | SRR19633495 | SRR19634229 |
| F022_E0E | SAMN29006945 | SRR19633484 | SRR19634218 |
| F022_EPA | SAMN29006946 | SRR19633473 | SRR19634207 |
| F022_EPB | SAMN29006947 | SRR19633462 | SRR19634196 |
| F022_EPC | SAMN29006948 | SRR19633451 | SRR19634185 |
| F022_EPD | SAMN29006949 | SRR19633445 | SRR19634179 |
| F022_EPE | SAMN29006950 | SRR19633444 | SRR19634178 |
| F022_EXA | SAMN29006951 | SRR19633516 | SRR19634250 |
| F022_EXB | SAMN29006952 | SRR19633515 | SRR19634249 |
| F022_EXC | SAMN29006953 | SRR19633514 | SRR19634248 |
| F022_EXD | SAMN29006954 | SRR19633513 | SRR19634247 |
| F022_EXE | SAMN29006955 | SRR19633512 | SRR19634246 |
| F054_E0A | SAMN29006956 | SRR19633511 | SRR19634245 |
| F054_E0B | SAMN29006957 | SRR19633510 | SRR19634244 |
| F054_E0C | SAMN29006958 | SRR19633509 | SRR19634243 |
| F054_E0D | SAMN29006959 | SRR19633508 | SRR19634242 |
| F054_E0E | SAMN29006960 | SRR19633507 | SRR19634241 |
| F054_EPA | SAMN29006961 | SRR19633505 | SRR19634239 |
| F054_EPB | SAMN29006962 | SRR19633504 | SRR19634238 |
| F054_EPC | SAMN29006963 | SRR19633503 | SRR19634237 |
| F054_EPD | SAMN29006964 | SRR19633502 | SRR19634236 |
| F054_EPE | SAMN29006965 | SRR19633501 | SRR19634235 |
| F054_EXA | SAMN29006966 | SRR19633500 | SRR19634234 |
| F054_EXB | SAMN29006967 | SRR19633499 | SRR19634233 |
| F054_EXC | SAMN29006968 | SRR19633498 | SRR19634232 |
| F054_EXD | SAMN29006969 | SRR19633497 | SRR19634231 |
| F054_EXE | SAMN29006970 | SRR19633496 | SRR19634230 |
| F104_E0A | SAMN29006971 | SRR19633494 | SRR19634228 |
| F104_E0B | SAMN29006972 | SRR19633493 | SRR19634227 |
| F104_E0C | SAMN29006973 | SRR19633492 | SRR19634226 |
| F104_E0D | SAMN29006974 | SRR19633491 | SRR19634225 |
| F104_E0E | SAMN29006975 | SRR19633490 | SRR19634224 |
| F104_EPA | SAMN29006976 | SRR19633489 | SRR19634223 |
| F104_EPB | SAMN29006977 | SRR19633488 | SRR19634222 |
| F104_EPC | SAMN29006978 | SRR19633487 | SRR19634221 |
| F104_EPD | SAMN29006979 | SRR19633486 | SRR19634220 |
| F104_EPE | SAMN29006980 | SRR19633485 | SRR19634219 |
| F104_EXA | SAMN29006981 | SRR19633483 | SRR19634217 |
| F104_EXB | SAMN29006982 | SRR19633482 | SRR19634216 |
| F104_EXC | SAMN29006983 | SRR19633481 | SRR19634215 |
| F104_EXD | SAMN29006984 | SRR19633480 | SRR19634214 |
| F104_EXE | SAMN29006985 | SRR19633479 | SRR19634213 |
| MG1655_E0A | SAMN29006986 | SRR19633478 | SRR19634212 |
| MG1655_E0B | SAMN29006987 | SRR19633477 | SRR19634211 |
| MG1655_E0C | SAMN29006988 | SRR19633476 | SRR19634210 |
| MG1655_E0D | SAMN29006989 | SRR19633475 | SRR19634209 |
| MG1655_E0E | SAMN29006990 | SRR19633474 | SRR19634208 |
| MG1655_EPA | SAMN29006991 | SRR19633472 | SRR19634206 |
| MG1655_EPB | SAMN29006992 | SRR19633471 | SRR19634205 |
| MG1655_EPC | SAMN29006993 | SRR19633470 | SRR19634204 |
| MG1655_EPD | SAMN29006994 | SRR19633469 | SRR19634203 |
| MG1655_EPE | SAMN29006995 | SRR19633468 | SRR19634202 |
| MG1655_EXA | SAMN29006996 | SRR19633467 | SRR19634201 |
| MG1655_EXB | SAMN29006997 | SRR19633466 | SRR19634200 |
| MG1655_EXC | SAMN29006998 | SRR19633465 | SRR19634199 |
| MG1655_EXD | SAMN29006999 | SRR19633464 | SRR19634198 |
| MG1655_EXE | SAMN29007000 | SRR19633463 | SRR19634197 |
| ELU39_E0A | SAMN29007001 | SRR19633461 | SRR19634195 |
| ELU39_E0B | SAMN29007002 | SRR19633460 | SRR19634194 |
| ELU39_E0C | SAMN29007003 | SRR19633459 | SRR19634193 |
| ELU39_E0D | SAMN29007004 | SRR19633458 | SRR19634192 |
| ELU39_E0E | SAMN29007005 | SRR19633457 | SRR19634191 |
| ELU39_EPA | SAMN29007006 | SRR19633456 | SRR19634190 |
| ELU39_EPB | SAMN29007007 | SRR19633455 | SRR19634189 |
| ELU39_EPC | SAMN29007008 | SRR19633454 | SRR19634188 |
| ELU39_EPD | SAMN29007009 | SRR19633453 | SRR19634187 |
| ELU39_EPE | SAMN29007010 | SRR19633452 | SRR19634186 |
| ELU39_EXA | SAMN29007011 | SRR19633450 | SRR19634184 |
| ELU39_EXB | SAMN29007012 | SRR19633449 | SRR19634183 |
| ELU39_EXC | SAMN29007013 | SRR19633448 | SRR19634182 |
| ELU39_EXD | SAMN29007014 | SRR19633447 | SRR19634181 |
| ELU39_EXE | SAMN29007015 | SRR19633446 | SRR19634180 |

**Supplementary Table 2.** Accession numbers for all MG1655 transcriptomic data. Samples were sequenced in triplicate, and include ancestral isolate without plasmid (A0), ancestral isolate with plasmid (AP), and experimentally evolved isolates with 5 evolution replicates (AE), and 3 RNAseq replicates (1-3). E0 = evolved without plasmid, EP = evolved with plasmid, EX = evolved with plasmid in the presence of cefotaxime.

| Isolate | Biosample | RNA Replicate 1 | RNA Replicate 2 | RNA Replicate 3 |
| --- | --- | --- | --- | --- |
| MG1655_A0 | SAMN29009610 | SRR19635002 | SRR19635000 | SRR19634998 |
| MG1655_AP | SAMN29009611 | SRR19635001 | SRR19634999 | SRR19634997 |
| MG1655_E0A | SAMN29006986 | SRR19646225 | SRR19646218 | SRR19646201 |
| MG1655_E0B | SAMN29006987 | SRR19646224 | SRR19646217 | SRR19646200 |
| MG1655_E0C | SAMN29006988 | SRR19646213 | SRR19646216 | SRR19646199 |
| MG1655_E0D | SAMN29006989 | SRR19646202 | SRR19646215 | SRR19646198 |
| MG1655_E0E | SAMN29006990 | SRR19646191 | SRR19646214 | SRR19646197 |
| MG1655_EPA | SAMN29006991 | SRR19646185 | SRR19646212 | SRR19646196 |
| MG1655_EPB | SAMN29006992 | SRR19646184 | SRR19646211 | SRR19646195 |
| MG1655_EPC | SAMN29006993 | SRR19646183 | SRR19646210 | SRR19646194 |
| MG1655_EPD | SAMN29006994 | SRR19646182 | SRR19646209 | SRR19646193 |
| MG1655_EPE | SAMN29006995 | SRR19646181 | SRR19646208 | SRR19646192 |
| MG1655_EXA | SAMN29006996 | SRR19646223 | SRR19646207 | SRR19646190 |
| MG1655_EXB | SAMN29006997 | SRR19646222 | SRR19646206 | SRR19646189 |
| MG1655_EXC | SAMN29006998 | SRR19646221 | SRR19646205 | SRR19646188 |
| MG1655_EXD | SAMN29006999 | SRR19646220 | SRR19646204 | SRR19646187 |
| MG1655_EXE | SAMN29007000 | SRR19646219 | SRR19646203 | SRR19646186 |

